# Multiscale confidence quantification for virtual spatial transcriptomics with UTOPIA

**DOI:** 10.64898/2026.03.01.708850

**Authors:** Kaitian Jin, Zihao Chen, Xiaokang Yu, Musu Yuan, Amelia Schroeder, Bernhard Dumoulin, Yunhe Liu, Linghua Wang, Jeong Hwan Park, Tae Hyun Hwang, Katalin Susztak, Zhimei Ren, Nancy R. Zhang, Mingyao Li

## Abstract

Virtual spatial transcriptomics (ST) methods predict gene expression or cell types from histology images, extending molecular readouts beyond the limited regions or samples directly measured by ST platforms. However, the statistical reliability of these predictions remains unclear. Here, we present UTOPIA, a model-agnostic framework for multiscale confidence quantification in virtual ST. UTOPIA assigns statistically calibrated confidence scores to predictions across spatial resolutions and biological granularities, ranging from single genes to metagenes and from specific cell types to broader cell classes. UTOPIA controls false discovery rates for detecting genes, metagenes, or cell types while accounting for local tissue context. We show that prediction confidence depends critically on both spatial resolution and biological granularity, with reliable inference often emerging only at coarser, biologically meaningful scales. Across multiple ST platforms and in both in-sample and out-of-sample settings, UTOPIA enhances interpretability, prevents false biological conclusions, and enables more trustworthy downstream analyses of virtual ST.

## Introduction

Spatial transcriptomics (ST) technologies provide spatially resolved measurements of gene expression within intact tissue sections, enabling inference of cell types and offering unprecedented insights into tissue organization, disease mechanism and microenvironments of tissues ^1-5^. Despite their promise, the high cost, long turnaround time, and limited capture area of current ST platforms constrain their widespread adoption in biomedical research and clinical practice.

Virtual ST provides a practical alternative to high-cost molecular profiling, where gene expression or cell types are inferred directly from hematoxylin and eosin (H&E)-stained histology images through models trained on coupled H&E and ST data ^6-10^. H&E staining is inexpensive and ubiquitous in pathology, capturing rich morphological information that can be predictive of underlying molecular states. Motivated by their low cost and high information content, a diverse array of computational models has been developed to predict molecular profiles from H&E images across scales, from low-resolution spots ^7, 11, 12^, to mixed resolution ^13^, and super-resolution ^14-19^. Typically, these approaches rely on ST measurements from selected, histologically representative regions of interest (ROIs), chosen either manually or automatically ^20^, to train prediction models that are then used to generate virtual ST across unmeasured regions of the tissue or other tissue samples.

Despite these advances, a critical barrier to the broader adoption of virtual ST is the lack of principled uncertainty quantification. Prediction accuracy varies widely across datasets, molecular targets, spatial locations, and spatial resolutions, yet predictions are often interpreted as uniformly reliable. A model that performs well on one dataset may generalize poorly to another, and even within a single tissue section, prediction performance can differ substantially across genes and cell types. Consequently, when presented with virtual ST outputs, researchers lack a systematic framework for determining which predictions are statistically supported and which may reflect model artifacts. As predictive models grow increasingly complex^21-23^, their expressive power is accompanied by greater risk of overconfident or unsupported inferences. To realize the potential of virtual ST prediction, rigorous and statistically grounded approaches to quantifying prediction uncertainty are urgently needed.

This challenge raises fundamental questions. First, which molecular targets can be predicted with confidence? Genes with low measurement quality in the training data or weak associations with histological features are generally challenging to predict; however, aggregating related genes into meta-genes can improve data quality and strengthen their associations with histological features. Similarly, models often struggle to distinguish morphologically similar cell types, such as B and T cells, solely based on histology; grouping them into more broadly defined cell classes, such as lymphocytes, can substantially improve prediction accuracy.

Second, at what spatial resolution should predictions be trusted? Although many models produce fine-grained or super-resolution outputs by default, many biological tasks, such as identifying tertiary lymphoid structures (TLS) ^24-27^, do not require such high resolution. Moreover, visual similarity between predicted and measured spatial patterns can be misleading, as comparisons are often performed at coarse visual scales while predictions are generated at much finer resolution. Assessing prediction confidence at biologically relevant spatial scales is therefore essential.

Third, at what expression magnitude do predictions meaningfully discriminate signal from noise? All current methods score low on standard evaluation metrics, such as Pearson correlation coefficient, for gene expression prediction. For example, a recent benchmarking study ^28^ found that the Pearson correlation between predicted and measured expression of genes averages around 0.2 for the best-performing methods. In many applications, however, precise expression prediction is unnecessary and it is sufficient to determine whether gene expression exceeds a biologically meaningful threshold. Indeed, prior work ^29^ has demonstrated that binarized expression is adequate for downstream tasks such as spatial clustering.

Finally, where within a tissue can predictions be trusted? Prediction accuracy varies spatially due to heterogeneity in tissue architecture, sample preservation, and the strength of local correlations between histological features and gene expression. Spatially resolved confidence estimates allow researchers to prioritize high-confidence regions for further investigation, guide targeted experimental validation, and avoid drawing conclusions from predictions in low-confidence areas.

Here we introduce UTOPIA (**U**ncertainty-aware **T**rustworthy Tools for Spatial **O**mics **P**redictions in **A**ll Scales), a general, model-agnostic framework for quantifying prediction confidence in virtual ST. UTOPIA systematically addresses the fundamental challenges of target identifiability, spatial resolution, expression magnitude, and spatial heterogeneity by assigning statistically calibrated confidence scores to predicted gene expression and cell types. The framework provides multiscale confidence quantification across two dimensions: spatial resolution, from coarse spot-level to super-resolution predictions, and biological granularity, from individual genes and cell types to functionally related meta-genes and broader cell classes. By guiding inference towards biologically meaningful and statistically robust scales, UTOPIA supports rigorous uncertainty quantification tailored to specific analytical tasks. We demonstrate the applicability of UTOPIA to both in-sample and out-of-sample predictions across multiple ST technologies, including Xenium, Xenium 5K, Visium HD, and CosMx. Using conformal inference, UTOPIA controls the false discovery rate (FDR) for detecting positive molecular signals and enhances downstream analyses, such as spatial clustering, compared to uncalibrated predictions. This provides a principled framework for trustworthy virtual spatial omics across diverse biological contexts.

## Results

### Overview of UTOPIA

UTOPIA addresses a fundamental challenge in virtual ST: quantifying confidence in predictions derived from histology images. In a typical in-sample virtual prediction workflow, a whole slide image (WSI) is available on a large H&E-stained tissue section, while molecular measurements are obtained only from smaller, information-rich regions of interest (ROIs) profiled using ST technologies (**Fig. 1**, left). Prediction models trained on these ROIs are then used to extrapolate gene expression or cell types across the unmeasured tissue. Although modern virtual ST models can generate predictions at super-resolution, approximately 8 μm × 8 μm and approaching single-cell resolution, these outputs are typically provided without reliability estimates, making it difficult to distinguish accurate predictions from artifacts.

**Fig. 1.**
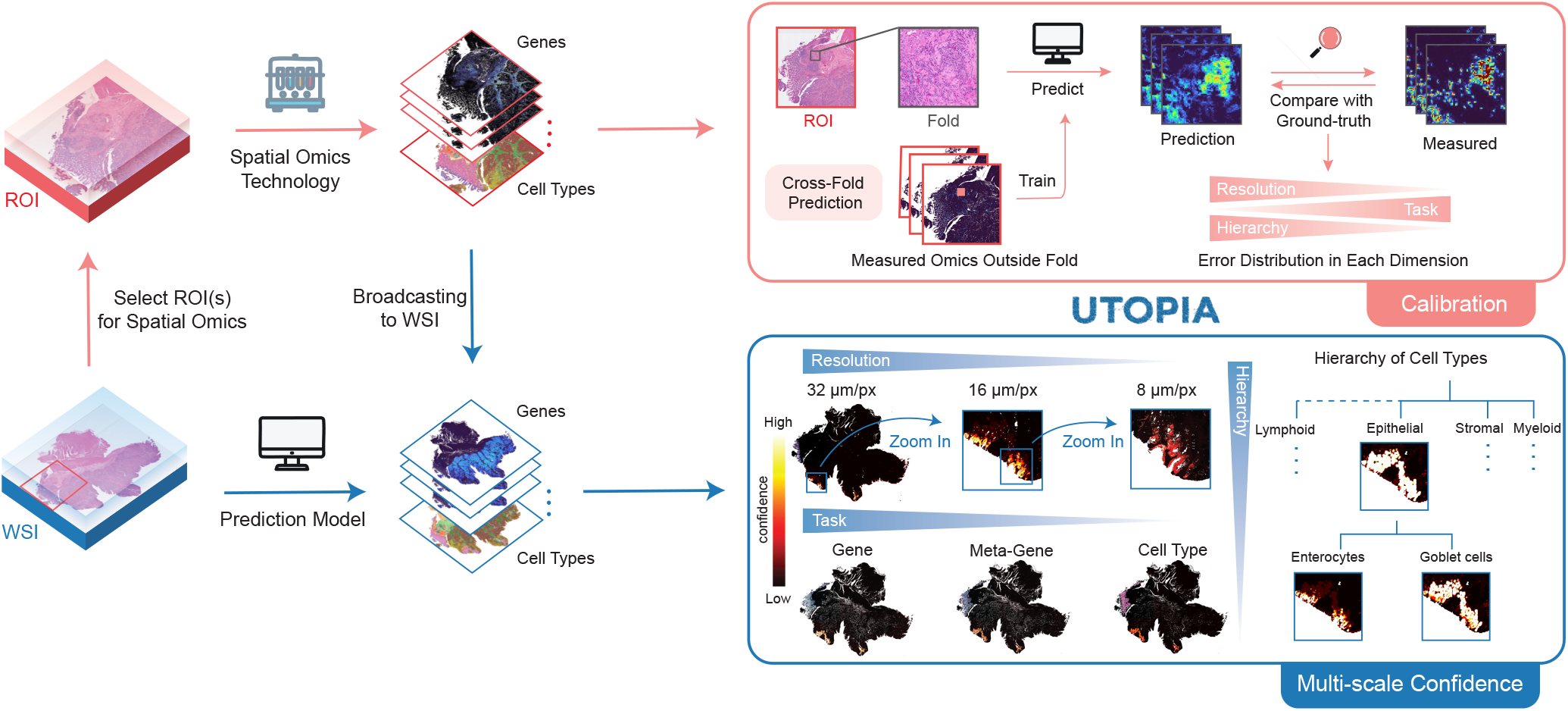
Overview of UTOPIA. Spatial omics data measured from regions of interest (ROIs) are used to train prediction models that can infer molecular information across whole slide images (WSI) (left panel). To calibrate prediction confidence, UTOPIA employs cross-fold validation by masking portions of the ROI for training and testing (upper right). Predictions are compared against ground-truth measurements to characterize error distributions across multiple dimensions. The calibrated error distributions enable UTOPIA to generate multi-scale confidence scores by conformal inference for predictions broadcast to the entire WSI (lower right).

UTOPIA provides statistically grounded confidence quantification for the outputs of any virtual prediction model (**Fig. 1**, right). The framework operates in two stages. First, UTOPIA constructs calibration data by partitioning ROIs into cross-validation folds that preserve histological diversity outside each fold (**Fig. 1**, top right). For each fold, a surrogate model is trained on the remaining folds and used to generate predictions on the held-out fold, thereby avoiding evaluation on the same data used for training. Comparing these held-out predictions with the corresponding observed measurements provides an empirical characterization of prediction errors across spatial resolutions and biological targets. In this calibration step, the empirically observed prediction errors in the held-out data are used to calibrate model-derived predictions that can be inaccurate.

In the second stage (**Fig. 1**, bottom right), UTOPIA leverages this calibrated error distribution to evaluate predictions in unmeasured regions of the tissue. For each spatial location and prediction target, UTOPIA identifies locations in calibration data with similar histological features and uses their empirically observed prediction errors to evaluate the model-derived prediction at the query location. This leverages the principle that histologically similar regions tend to exhibit similar observed molecular profiles and predictions thereof, leading to similar prediction uncertainty. Using conformal inference, UTOPIA computes a p-value for each spatial unit (a high-resolution pixel or a low-resolution patch) and each prediction target, testing the null hypothesis that the target is absent in the spatial unit. After multiple testing correction, these p-values are converted to confidence scores that reflect a calibrated FDR. Users can then identify spatial regions where gene expression or cell type presence is supported at a specified FDR level.

UTOPIA is model-agnostic and can be applied to predictions generated by any virtual ST method. Although we demonstrate the performance of UTOPIA using a simple neural network-based virtual ST approach, UTOPIA can be readily applied to more complex prediction models. In addition to detecting positive gene expression, UTOPIA supports confidence quantification for whether the predicted expression levels or cell type abundances exceed any user-specified threshold, which enables flexible applications across diverse biological settings. The combination of flexibility and principled uncertainty quantification makes UTOPIA a general-purpose tool for statistically rigorous and trustworthy interpretation of virtual ST predictions.

### Application to a human gastric cancer sample to predict tertiary lymphoid structures

To demonstrate how UTOPIA addresses the four questions outlined in the Introduction pertaining to target identification, spatial resolution, expression magnitude, and spatial heterogeneity, we applied UTOPIA to a large human gastric cancer tissue sample. This application illustrates how confidence quantification enables principled navigation of the trade-offs between prediction resolution and reliability, ultimately guiding interpretation at biologically appropriate scales.

We analyzed a 12 mm × 24 mm gastric cancer tissue section with matched H&E image (**Fig. 2A**, left and **Extended Data Fig. 1A**, left) and ST data using the 10X Genomics Xenium platform, profiling 377 genes and 14 manually annotated cell types (**Fig. 2A**, right and **Extended Data Fig. 1A**, right). Following the in-sample prediction workflow described in **Fig. 1**, we selected two 4 mm × 4 mm ROIs (red boxes in **Fig. 2A**) based on histological segmentation of 15 clusters (**Fig. 2A**, center and **Extended Data Fig. 1A**, center). These ROIs were chosen to capture all histology clusters present in the WSI to maximize their representativeness. A feedforward neural network (see **Methods**) was trained to learn the relationship between histological features extracted by the UNI foundation model ^30^ and molecular profiles within the ROIs and was then applied to predict molecular information across the entire unmeasured tissue.

**Fig. 2.**
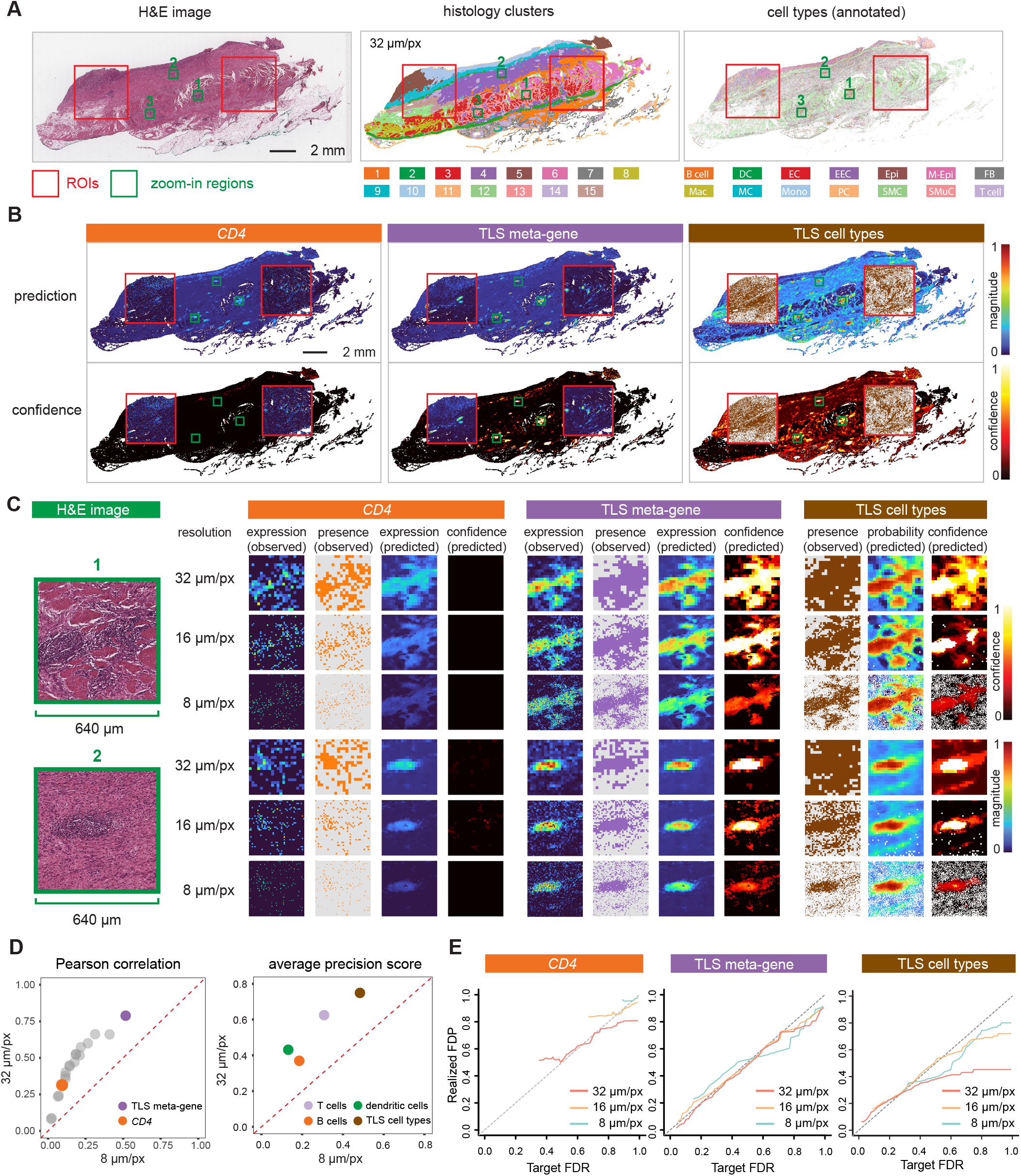
Application to a human gastric cancer sample to predict TLSs. **A**, The H&E image, histology clusters, and Xenium-based cell type annotation. Two ROIs (red) were selected for in-sample prediction. Cell type abbreviations: DC, dendritic cells; EC, endothelial cells; EEC, enteroendocrine cells; Epi, epithelial cells; M-Epi, malignant epithelial cells; FB, fibroblasts; Mac, macrophages; MC, mast cells; Mono, monocytes; PC, plasma cells; SMC, smooth muscle cells; SMuC, surface mucous cells. **B**, predictions and confidence scores at 32 µm/pixel resolution. For *CD4* and TLS meta-gene, predicted magnitude values are shown alongside UTOPIA-derived confidence scores. For TLS-associated cell types, softmax probability scores and corresponding confidence values are displayed. Confidence scores quantify the reliability of detecting the presence of each molecular feature. Ground-truth was displayed inside the ROIs. **C**, Multi-resolution analysis of zoom-in regions across 8, 16, and 32 µm/pixel resolutions. For *CD4* and TLS meta-gene: columns show ground-truth expression magnitude, binarized ground truth presence, predicted expression magnitude, and confidence for presence detection. For TLS cell types: ground-truth cell type annotations, softmax probability scores, and confidence for presence. **D**, Prediction performance metrics. Left: Pearson correlation coefficients between predicted and measured values at 8 and 32 µm/pixel for the TLS meta-gene and its constituent genes. *CD4* was highlighted in orange; gray points represent other genes within the TLS meta-gene signature. Right: Average precision scores for TLS cell type predictions at 8 and 32 µm/pixel resolution. **E**, FDR control using UTOPIA confidence scores. Portions of curves with fewer than 1% of total pixels meeting discovery criteria were omitted for clarity.

We focused our analysis on tertiary lymphoid structures (TLSs), organized lymphocyte aggregates with clinical relevance in cancer immunotherapy ^26, 27^. The presence of TLSs has often been associated with enhanced anti-tumor immunity and improved patient prognosis across multiple cancer types ^26, 27^, making its spatial mapping clinically important. TLSs comprise aggregates of B cells, T cells, and dendritic cells, each of which can be detected through specific marker expression, coordinated gene programs, and spatial co-localization of key immune lineages. This setting represents a common challenge for virtual ST, as individual marker genes, such as *CD4*, are often expressed sparsely and difficult to predict. In contrast, functionally related gene sets, such as a meta-gene formed by all TLS markers, may provide more robust prediction targets.

**Fig. 2B** shows predictions at the 32 μm per pixel resolution for three complementary representations of a TLS at different biological granularity: a single T cell marker gene *CD4*, a curated TLS meta-gene (**Extended Data Table 1**), and TLS aggregated cell types consisting of B, T, and dendritic cells. Although the prediction of all three targets displays broadly similar spatial patterns, UTOPIA’s confidence scores reveal striking differences in reliability. At the 32 μm resolution, UTOPIA assigned low confidence to *CD4* detection across the entire slide, but high confidence for predictions of both TLS meta-gene and TLS aggregated cell types. These results demonstrate that the granularity of biological representation matters: *CD4* cannot be reliably predicted as an individual gene, but combining functionally related genes or cell types substantially improved prediction confidence. Because our objective is to detect TLSs as organized immune structures rather than to quantify any single marker gene, inference at a coarser level of biological granularity is both sufficient and statistically reliable.

Next, we ask what the optimal spatial resolution for reliable prediction is. Consider three TLS-enriched regions (green boxes in **Fig. 2A, 2B**) at progressively finer resolutions of 32 μm, 16 μm, and 8 μm per pixel. As shown in **Fig. 2C** and **Extended Data Fig. 2B**, confidence scores decreased with increasing spatial resolution. At 8 μm, Xenium-detected gene expression is sparsely distributed across cells, whereas model predictions are spatially smooth, leading to reduced coherence with observed measurements and thus, reduced confidence. In contrast, the TLS meta-gene and aggregated cell types can be predicted at high confidence at the resolutions of 32 μm and 16 μm, with predictions aligning well with the observed presence of molecular signal. We have seen the trend of decreasing confidence along with increasing resolution beyond these three regions in the whole slide (**Extended Data Figs. 1D** and **2A**).

We validated UTOPIA’s confidence assessments by comparing prediction quality metrics across resolutions. For all genes in the TLS meta-gene, predictions at the 32 μm resolution showed higher Pearson correlation coefficient (PCC) with Xenium measurements than predictions at the 8 μm resolution (**Fig. 2D**, left). TLS meta-gene predictions were consistently more reliable than those for individual genes, across all resolutions. Similarly, predictions for the TLS aggregated cell types and individual cell types yielded a higher average precision score at the 32 μm resolution compared to the 8 μm resolution (**Fig. 2D**, right). The TLS aggregated cell types achieved even higher scores compared to individual cell types. These results confirm that UTOPIA’s resolution-dependent and target-dependent confidence estimates accurately reflect underlying prediction quality.

UTOPIA further provides rigorous FDR control (**Fig. 2E**). For TLS meta-gene, the empirical FDRs closely track the nominal levels across multiple target FDR thresholds, validating the statistical guarantees of conformal inference. In contrast, *CD4* has barely any discoveries when the target FDR is less than 50%, indicating that it is hard to predict the presence of *CD4* without making many errors. One reason is that Xenium’s capture efficiency for individual genes introduces measurement noise and thus lowers confidence. In contrast, the meta-gene that is derived from multiple genes exhibits superior measurement quality and controls false discoveries across all FDR thresholds. Indeed, UTOPIA achieved FDR control for additional pathway-level predictions, including active tumor angiogenesis, metastatic phenotype, and complement-mediated tumor inflammation (**Extended Data Fig. 2C** and **Extended Data Table 1**). These results highlight that training data quality is a key determinant of prediction confidence, a relationship we will explore systematically in subsequent experiments.

This case study demonstrates how UTOPIA guides researchers to identify the appropriate spatial resolution and granularity for trustworthy predictions. Rather than defaulting to single-marker predictions at the highest possible spatial resolution, researchers can use UTOPIA to determine the combinations of spatial resolution and biological granularity that yield both confidence and interpretability. In the example of TLS detection, reasonable confidence can be achieved at the intermediate resolution of 32 μm per pixel, using meta-genes or aggregated cell types. Super-resolution or single-gene predictions are neither reliable nor necessary.

### UTOPIA resolves spatially distinct cancer-associated fibroblast subtypes and their interactions with cancer cell states

Next, we applied UTOPIA to characterize cancer-associated fibroblast (CAF) heterogeneity in human cervical and ovarian cancer samples profiled using Xenium Prime 5K Human Pan Tissue & Pathways Panel (**Fig. 3A**). CAFs represent a critical component of the tumor microenvironment, and recent studies have identified functionally distinct CAF subtypes with unique spatial localization and cellular interaction ^31^. Understanding the distribution of CAF subtypes in situ is essential for deciphering tumor-stroma interactions and predicting therapeutic outcomes.

**Fig. 3.**
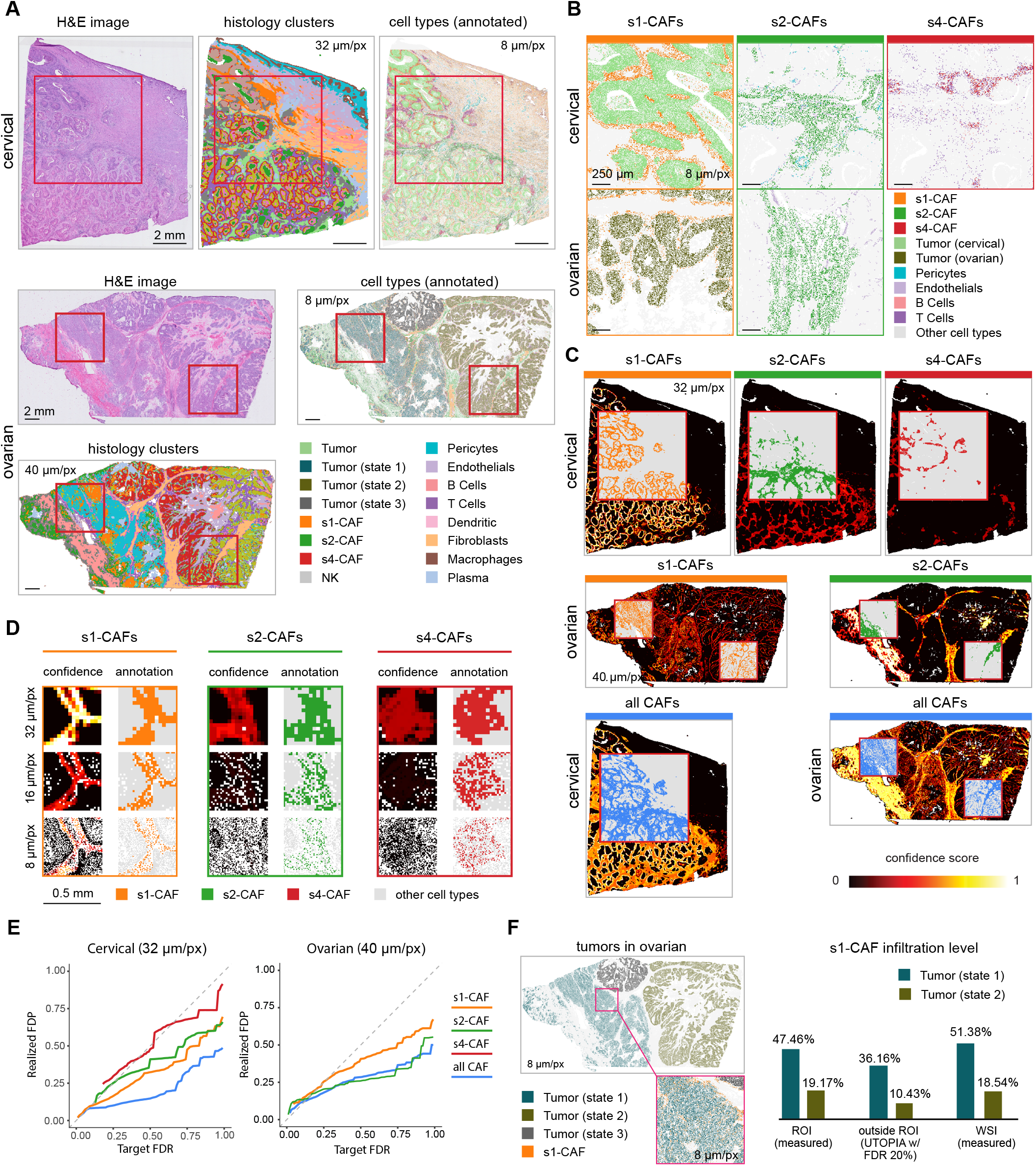
Application to investigate cancer-associated fibroblasts (CAFs) and their interactions with tumors. **A**, The H&E images, histology clusters, and annotated cell types of the cervical and ovarian sample. An ROI of size 6.4 mm × 6.4 mm was selected for the cervical sample. Two ROIs of size 4 mm × 4mm were selected for the ovarian sample. Cell types were annotated at 8 µm per pixel. Histology clusters were segmented at 32 and 40 µm per pixel for the two samples. **B**, Illustration of three different subtypes of CAFs in the two samples. Annotated cell types were displayed. **C**, Confidence scores for s1, s2, and s4-CAFs, as well as the combination of these three CAFs in the two samples. Confidence was evaluated in terms of the presence of corresponding molecules. Annotation of the cell type was displayed inside the ROIs. **D**, Three zoomed-in regions outside the ROI in the cervical sample. Each region is of size 0.5 mm × 0.5 mm with a focus on a subtype of CAFs. The highlighted pixels in the ground-truth are areas that contain at least one such cell. **E**, FDR control using UTOPIA confidence scores. Portions of curves with fewer than 1% of total pixels meeting discovery criteria were omitted for clarity. **F**, Infiltration levels of s1-CAF in tumors. Left: distribution of tumor cells and a zoomed-in window showing s1-CAF infiltration in state 1 tumor. Right: percentages of tumor cells that are infiltrated by s1-CAF.

Building on prior work that characterized four CAF subtypes (s1, s2, s3, and s4-CAFs) emerging from distinct cellular neighborhoods, we sought to detect these cell populations across entire tissue sections. Notably, s1-CAFs localize predominantly adjacent to cancer cells, s2-CAFs largely occupy stromal regions, and s4-CAFs associate with B and T cell aggregates (**Fig. 3B**). The s3-CAF subtype was absent from both samples examined. The intermingling of s4-CAFs with lymphocytes, contrasted with the discrete localization of s1-CAFs near cancer cells, creates varying levels of prediction difficulty that UTOPIA can quantify.

We selected a 6.4 mm × 6.4 mm ROI from the cervical cancer sample and two 4 mm × 4 mm ROIs from the ovarian cancer sample to capture histological diversity representative of the whole slide (**Fig. 3A**). Cell type annotation within the cervical ROI identified s1, s2, and s4-CAFs, while the ovarian ROIs contained s1 and s2-CAFs (**Extended Data Fig. 3A**). Following the in-sample virtual prediction workflow, we generated softmax probability scores for all CAF subtypes across unmeasured tissue regions (**Extended Data Figs. 3B** and **3C**), then applied UTOPIA to quantify confidence for their presence at resolutions of 32 μm (cervical) and 40 μm (ovarian) per pixel.

At these resolutions, UTOPIA’s confidence scores revealed striking differences in prediction reliability across CAF subtypes (**Fig. 3C**). UTOPIA shows high confidence for the presence of s1-CAFs and low confidence for that of s4-CAFs. Examining finer resolutions at one zoom-in region (**Fig. 3D**, left) on the cervical cancer sample demonstrates that confident detection of s1-CAF presence remains achievable even at 8 μm per pixel. This high confidence stems from s1-CAFs’ clear spatial segregation adjacent to cancer cells without intermingling with other cell populations (**Fig. 3B**). In contrast, s4-CAFs spatially intermingle with B and T cells, creating morphological ambiguity that inherently limits how the virtual ST model can predict their presence based on histological features alone. UTOPIA’s confidence scores successfully captured this biological reality: subtypes with distinct spatial niches yield high confidence in their predicted presence, while subtypes embedded within heterogeneous cellular neighborhoods generate appropriately lower confidence assessments. Quantitatively, UTOPIA maintains rigorous FDR control across both samples and all CAF subtypes (**Fig. 3E**). Importantly, this analysis demonstrates that coarser resolutions (40 μm/px) suffice for confidently identifying the presence of distinct cell type populations with characteristic spatial localization. The confidence scores for s1-CAF and s2-CAF presence at 32 μm/px on the cervical sample display clearly distinct spatial patterns (**Fig. 3C** and **3D**), enabling researchers to identify biologically relevant tumor microenvironments without requiring super-resolution predictions.

Beyond quantifying confidence for CAF subtypes, we investigated whether UTOPIA could identify functionally relevant cancer cell-CAF interactions. Previous work ^31^ identified three cancer cell states in ovarian samples distinguished by differential s1-CAF infiltration levels. Among the three, state 1 cancer cells associate with high s1-CAF infiltration, while state 2 cancer cells exhibit minimal s1-CAF proximity (**Fig. 3F**, left). We first validated these associations within the profiled ROI (**Fig. 3F**, right), confirming the expected infiltration patterns with 47.46% of 40-μm bins containing state 1 tumor cells also contain s1-CAF while only 19.17% of those containing state 2 tumor cells contain s1-CAF. We then applied UTOPIA to quantify confidence for the presence of state 1 and state 2 cancer cells across unmeasured tissue, selecting 40-μm bins with confidence scores corresponding to FDR ≤ 20% for s1-CAFs, state 1 cancer cells, and state 2 cancer cells. Calculating the spatial overlap between high-confidence s1-CAF bins and high-confidence cancer cell state bins revealed that infiltration patterns in predicted regions closely recapitulate those observed within the ROI. This concordance validates that UTOPIA’s confidence assessments accurately reflect underlying biological relationships; regions confidently predicted to contain state 1 cancer cells indeed show elevated s1-CAF co-occurrence, while regions confidently predicted to contain state 2 cancer cells show reduced s1-CAF association.

### UTOPIA enables cross-platform generalization and reveals glomerular pathology in diabetic nephropathy

To validate the generalizability of UTOPIA across ST platforms and to assess its ability to reveal disease-associated tissue structural changes, we applied the framework to human kidney samples profiled using NanoString’s CosMx platform. CosMx differs fundamentally from the Xenium platform used in the previous examples^32-34^. Instead of profiling large contiguous ROIs, CosMx measures transcriptomic signals within preselected, small 0.5 mm × 0.5 mm field-of-views (FOVs). Additionally, CosMx exhibits lower molecule capture rates than Xenium, resulting in sparser cell type annotations as illustrated by the zoom-in views of FOVs 13 and 17 in **Fig. 4A** and additional FOVs in **Extended Data Fig. 6B**. This challenging prediction setting provides an informative test case for UTOPIA’s ability to quantify prediction confidence under suboptimal data conditions.

**Fig. 4.**
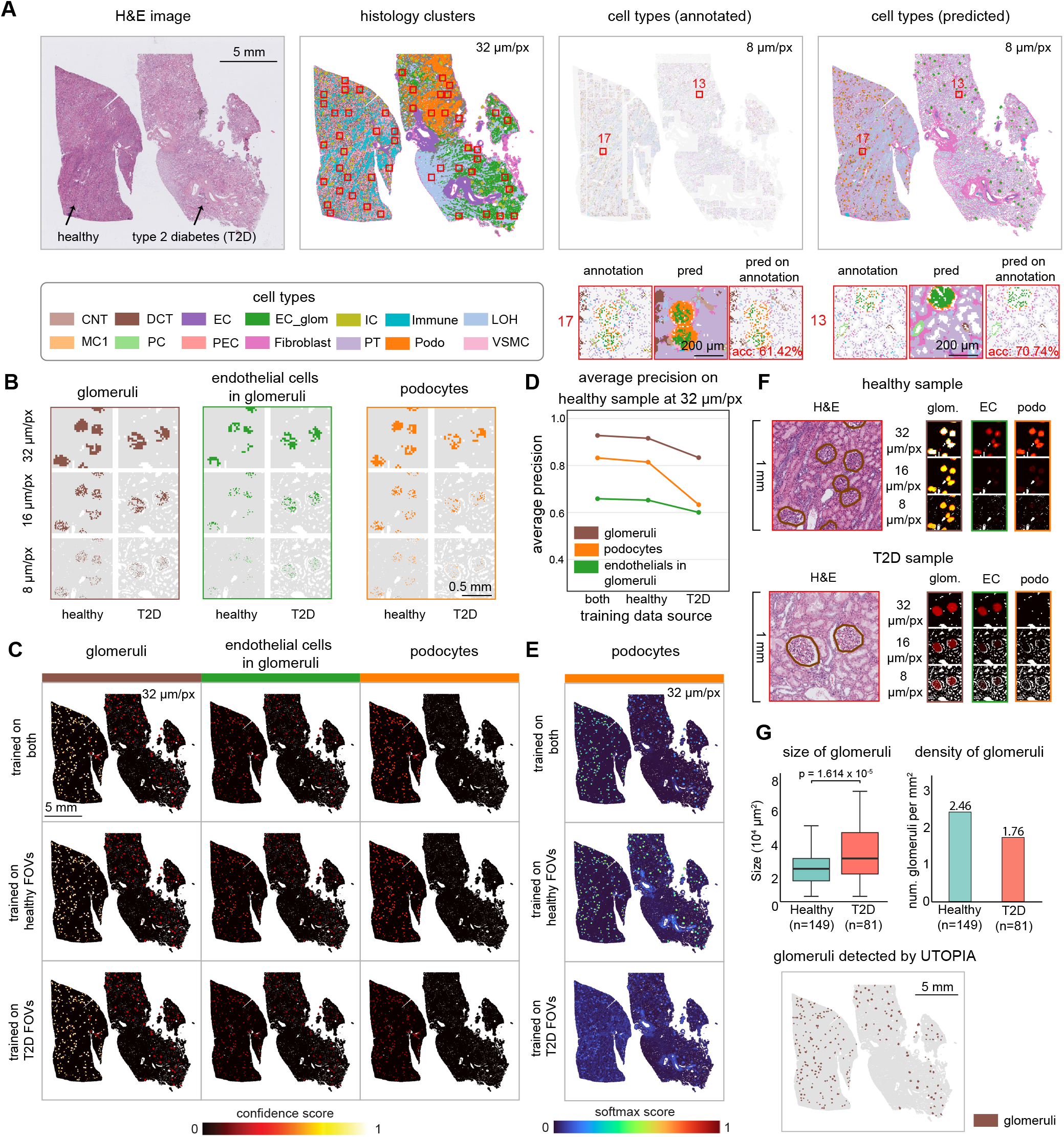
Detection of glomeruli in human kidney samples. **A**, The H&E images, histology clusters (32 µm/pixel), CosMx-based annotated cell types (8 µm/pixel), and predicted cell types (8 µm/pixel) of the healthy sample and the type 2 diabetes (T2D) sample. We selected 25 FOVs of size 0.5 mm × 0.5 mm for each sample. Annotated cell types inside the 50 FOVs were used to train a model to predict cell types outside the FOVs and uncaptured locations inside the FOVs. Two FOVs were zoomed in for illustrations. **B**, Zoomed-in visualization of annotated cell types in the two samples across 8, 16, and 32 µm/pixel resolutions. **C**, Confidence scores on the presence of cell types at 32 µm/pixel. Columns correspond to cell types. Rows are models trained on different FOVs. **D**, Average precision scores for three cell types on the healthy sample, when trained on different FOVs. **E**, Predicted softmax probability scores on podocytes at 32 µm/pixel. **F**, Zoomed-in windows for confidence scores at one area outside FOVs for each sample. **E**, Size and density of glomeruli detected under 32 µm/pixel on the two samples. Glomeruli were detected by first filtering 32-µm bins with confidence scores higher than 0.41 (corresponding to FDR at 10%) and then using connected component analysis.

We analyzed a comparative dataset in which kidney samples from two individuals were placed on the same slide and profiled simultaneously (**Fig. 4A**, left). The left sample was obtained from a healthy donor, whereas the right sample originated from a patient with type 2 diabetes (T2D). For each sample, we selected 25 0.5 mm × 0.5 mm FOVs denoted by red rectangles in **Fig. 4A** with further details on each FOV shown in **Extended Figs. 4 and 5**. Prediction models were trained using molecular information within these FOVs and used to predict cell types across unmeasured tissue regions as well as unannotated pixels within FOVs in both samples (**Fig. 4A**, right). As shown in **Fig. 4A** for FOVs 13 and 17 and additional FOVs in **Extended Data Fig. 6B**, virtual prediction enriched the sparsely annotated cell type labels in the FOVs but also had low accuracy (31.14% - 72.88%) at 8 μm resolution when we compared predictions with annotation, calling the importance of using UTOPIA to quantify the prediction uncertainty.

The design of virtual prediction on samples from two patients raises a fundamental question regarding training data composition: when predicting cell types in healthy tissue, should models be trained using all 50 FOVs from both samples or only the 25 FOVs from the healthy sample? While including all FOVs increases the number of cells in the training data, disease-associated inter-individual heterogeneity may compromise prediction reliability. UTOPIA provides a principled framework for evaluating this trade-off by quantifying prediction confidence under alternative training scenarios (see **Methods**).

We focused our analysis on glomeruli, the kidney’s filtration units that are centrally involved in diabetic nephropathy. Each glomerulus consists of a specialized capillary network comprised of three key cell types: glomerular endothelial cells (EC glom), podocytes (podo), and parietal epithelial cells (PEC). Due to the low molecular capture rate of CosMx, only a small number of 8-μm bins per glomerulus received cell type annotations, making glomerular structures difficult to resolve at high spatial resolution, such as 8 μm per pixel, as shown in **Fig. 4B**. However, when annotations are aggregated to coarser resolutions of 32 μm per pixel, glomerular architecture becomes readily evident (**Fig. 4B**) as well as podocytes and endothelial cells within glomeruli. We therefore evaluated UTOPIA’s confidence scores across multiple resolutions, including 32 μm, 16 μm, and 8 μm per pixel, for detecting glomeruli as composite structures as well as for individual glomerular endothelial cells and podocytes.

**Fig. 4C** shows confidence maps at 32 μm per pixel for three models trained on different subsets of FOVs (both samples, healthy sample only, and T2D sample only). For glomerular structure detection (**Fig. 4C**, first column), confidence scores were largely consistent across the three models, indicating that detecting glomeruli as composite units is robust to variations in training data composition.

For individual glomerular cell types, however, training data composition matters. For example, using all 50 FOVs from both samples did not substantially reduce prediction confidence for podocytes on the healthy sample compared to using only the 25 healthy FOVs (first vs. second row in the third column, **Fig. 4C**), suggesting that incorporating additional data from the T2D sample did not introduce meaningful heterogeneity. However, training exclusively on T2D FOVs (third row in the third columns, **Fig. 4C**) exhibited substantially lower confidence, corroborated by a corresponding decrease in average precision (**Fig. 4D**). This result highlights a critical caution that models trained on data from a different individual should be applied carefully, as inter-individual heterogeneity such as the well-documented reduction in podocyte density in T2D ^35-37^ can compromise prediction reliability.

Encouragingly, UTOPIA provides a safeguard against such misuse. The model trained exclusively on healthy FOVs assigned high predicted probabilities for podocytes in the T2D sample (**Fig. 4E**), where podocyte density is known to be reduced, a clear case of false positives driven by training data mismatch. UTOPIA successfully identified and corrected these false discoveries (third columns in **Fig. 4C**, FDR control curves in **Extended Fig. 8**), assigning low confidence to these predictions and demonstrating its ability to protect against misleading conclusions even when training data is poorly matched to the target sample. We further demonstrate a use case of UTOPIA in correcting false discoveries in the following section.

To examine spatial resolution effects in greater detail, we inspected higher-resolution predictions in two representative regions from each sample (**Fig. 4F**). At the 32 μm resolution, high-confidence regions successfully delineated the glomerular contours visible in the corresponding H&E images. As the resolution increased, confidence scores decreased systematically for individual cell types, reflecting the limited data quality within FOVs and the resulting difficulty in making reliable super-resolution predictions (confidence on the whole slide for 16 μm and 8 μm per pixel in **Extended Fig. 7**).

Visual inspection of the H&E images and the confidence-guided spatial maps revealed that glomeruli in the T2D sample appeared consistently larger than those in the healthy sample, suggesting glomerular hypertrophy in diabetic nephropathy. To quantify this observation, we selected 32 μm resolution super-pixels with high UTOPIA confidence scores for glomerular presence, corresponding to FDR ≤ 20%, and applied connected component analysis to segment individual glomeruli as shown in **Fig. 4G** (see **Methods**). Comparison of glomerular sizes across sections revealed a significant size increase in the T2D sample relative to the healthy sample (p =1.614 × 10^-5^, two-sample t-test; **Fig. 4G**, top left). Additionally, the healthy sample exhibited higher glomerular density at 2.46 glomeruli per mm^2^ (**Fig. 4G**, top right), consistent with the progressive glomerular loss that occurs in diabetic nephropathy as individual glomeruli undergo sclerosis^37^.

These findings demonstrate that UTOPIA can generate biologically meaningful spatial maps of prediction confidence even in challenging settings characterized by sparse measurements and platform-specific constraints. Beyond quantifying prediction reliability, the framework also enables detection of clinically relevant pathological changes, including podocyte depletion, glomerular hypertrophy and reduced glomerular density, that distinguish diabetic from healthy kidney tissue. Moreover, this analysis highlights UTOPIA’s utility in cross-individual prediction scenarios, where the representativeness of training data is often uncertain. Rather than requiring users to carefully curate training data upfront, UTOPIA provides a principled means to detect when training data mismatch leads to unreliable predictions, allowing users to be more relaxed about training data selection while maintaining statistical rigor.

### False positive control for out-of-subject predictions on normal gastric samples

To evaluate UTOPIA’s performance beyond in-sample prediction, we conducted out-of-subject, out-of-sample predictions on normal gastric samples. The training sample (N1) and test sample (N2) were obtained from different individuals and profiled using the Xenium platform with 377 genes (**Fig. 5A**). Joint clustering of post-Xenium H&E images from both sections identified 15 distinct histological clusters (**Fig. 5A**). The training sample served a dual role by providing data for both model training and calibration, analogous to the use of ROIs in the in-sample scenario. Thus, the test sample (N2) was masked during both the training and calibration step.

**Fig. 5.**
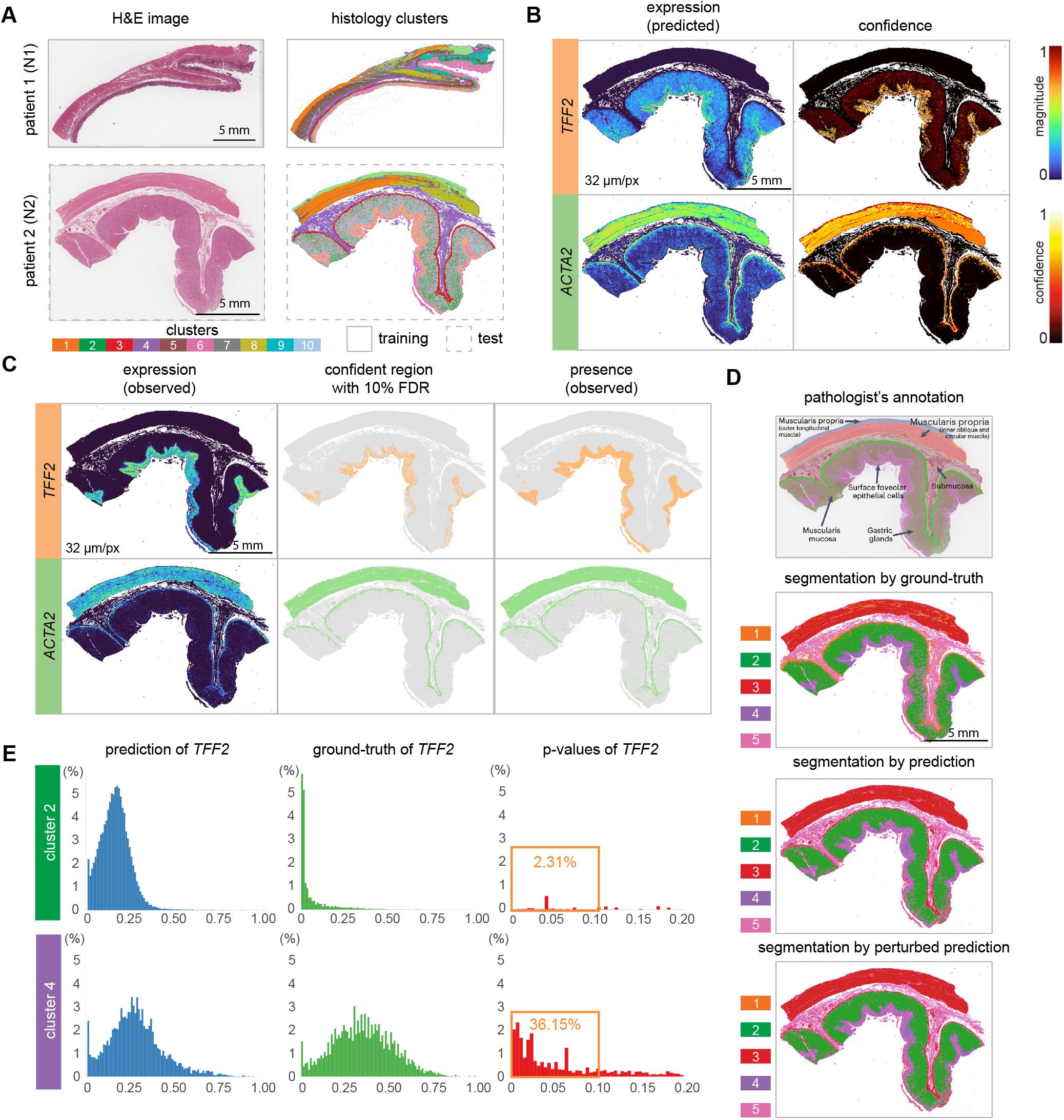
Correcting false positives in out-of-subject prediction. **A**, The H&E images and histology clusters (32 µm/pixel) of gastric tissue samples from two subjects. **B**, Predicted expression and confidence scores for *TFF2* and *ACTA2* at 32 µm/pixel. **C**, The first column is the predicted expression magnitude. The second column highlights 32-µm bins with confidence scores higher than 0.41 (corresponding to 10% FDR). The third column highlights 32-µm bins with expression of the two genes in the ground-truth. **D**, Segmentation of sample N2 based on different inputs. All but the pathologist’s annotation were done at 32 µm/pixel. **E**, Distribution of prediction, ground-truth, and UTOPIA’s p-values for *TFF2* in clusters 2 and 4 of the segmentation by prediction. The horizontal axis stands for magnitudes or p-values. The vertical axis stands for the percentages of 32-µm bins in the cluster. Bars in histograms higher than 6% were truncated at 6%.

Out-of-subject predictions on the test sample at 32 µm spatial resolution for two genes, *TFF2* and *ACTA2*, are shown in **Fig. 5B**. In the raw predictions, *TFF2* showed moderate expression within gastric glands, and *ACTA2* appeared sporadically in surface foveolar epithelium (pathology annotation in **Fig. 5D**). Regions with low predicted expression magnitudes introduced interpretive ambiguity. In practice, the absence of ground-truth measurements would make it difficult to distinguish whether such signals represent genuine low-level expression or prediction artifacts.

We applied UTOPIA to make the distinction. UTOPIA yielded interpretable confidence scores that distinguish supported from unsupported signals. As shown in **Fig. 5B**, UTOPIA assigned low confidence scores to the regions with low predicted expression magnitudes. To validate UTOPIA’s results, we next compared with the ground truth measurements (**Fig. 5C**, left), masked during training, which revealed that these less confident regions are indeed prediction artifacts. As shown in **Fig. 5C**, regions with high confidence scores (equivalent to 10% FDR) correspond to true positive regions, effectively separating artifactual from biologically supported predictions.

Virtual ST predictions are often used as inputs for downstream analyses such as tissue segmentation. Notably, segmentation based on predicted gene expression appeared comparable to pathologist annotation even in the presence of false positive signals (**Fig. 5D**). Simple segmentation methods such as K-means clustering were robust to moderate prediction errors; replacing 10% of gene expression predictions with random values drawn uniformly between 0 and 1 had minimal impact on segmentation results. This robustness might suggest that correcting individual prediction errors is unnecessary. However, while errors may not distort the segmentation itself, they can propagate into the biological interpretation of the resulting segments.

In the segmentation obtained from predicted expression, cluster 2 corresponding to gastric glands exhibited modest *TFF2* expression, while cluster 4 corresponding to surface foveolar epithelium showed modest-to-high *TFF2* expression (**Fig. 5E**, left). This pattern is biologically problematic as *TFF2* expression in normal gastric mucosa is primarily restricted to mucous neck cells in the isthmus region and is not characteristic of chief cells within the deeper gastric glands. Erroneous *TFF2* expression in gastric glands (cluster 2) could therefore be misinterpreted as early spasmolytic polypeptide-expressing metaplasia, a precancerous condition marked by transdifferentiate of chief cells into *TFF2*-expressing mucous cells.

UTOPIA prevented such misinterpretation by providing statistical evidence for or against gene expression in specific tissue regions (**Fig. 5E**, right). Among the 32-µm super-pixels within cluster 2, only 2.31% showed significant *TFF2* expression (adjusted P < 0.1), compared to 36.15% within cluster 4. Thus, thresholding by UTOPIA p-values correctly rejected the spurious expression of *TFF2* caused by inaccurate model prediction, preventing false biological conclusions in out-of-subject virtual ST analyses.

### Quality of training data affects confidence scores in UTOPIA

Across all prediction experiments, a consistent pattern has emerged: UTOPIA’s confidence scores strongly depend on the quality of the ST data used for training and calibration. Multiple factors contribute to variation in data quality. Individual genes often exhibit sparse molecular detection, whereas meta-genes demonstrate reduced noise and higher molecule counts. ST platforms also differ substantially in molecular capture efficiency; for example, imaging-based platforms typically detect more molecules per gene than sequencing-based platforms. Additionally, the quality of sample preservation affects measurement reliability. These sources of heterogeneity collectively determine the scale at which confident predictions can be achieved.

To systematically examine how data quality affects prediction confidence, as estimated by UTOPIA, we designed an experiment using a human lung adenocarcinoma sample profiled at multiple quality levels. This design leveraged a unique dataset in which the same tissue block was profiled three times across two platforms (**Fig. 6A**). One slide (top, red) underwent sequential profiling on Xenium followed by Visium HD, whereas a second slide (bottom, blue) from the same tissue block was profiled using Visium HD alone. After registering all three datasets to the H&E image of the red slide, we constructed three training datasets with varying quality: a high-quality dataset combining all three profiles, a medium-quality dataset combining the two Visium HD profiles, and a low-quality dataset consisting of the single Visium HD profile from the blue slide. All datasets included genes shared between the Xenium and Visium HD panels. We restricted analysis to the region profiled by all three ST datasets (**Fig. 6B**).

**Fig. 6.**
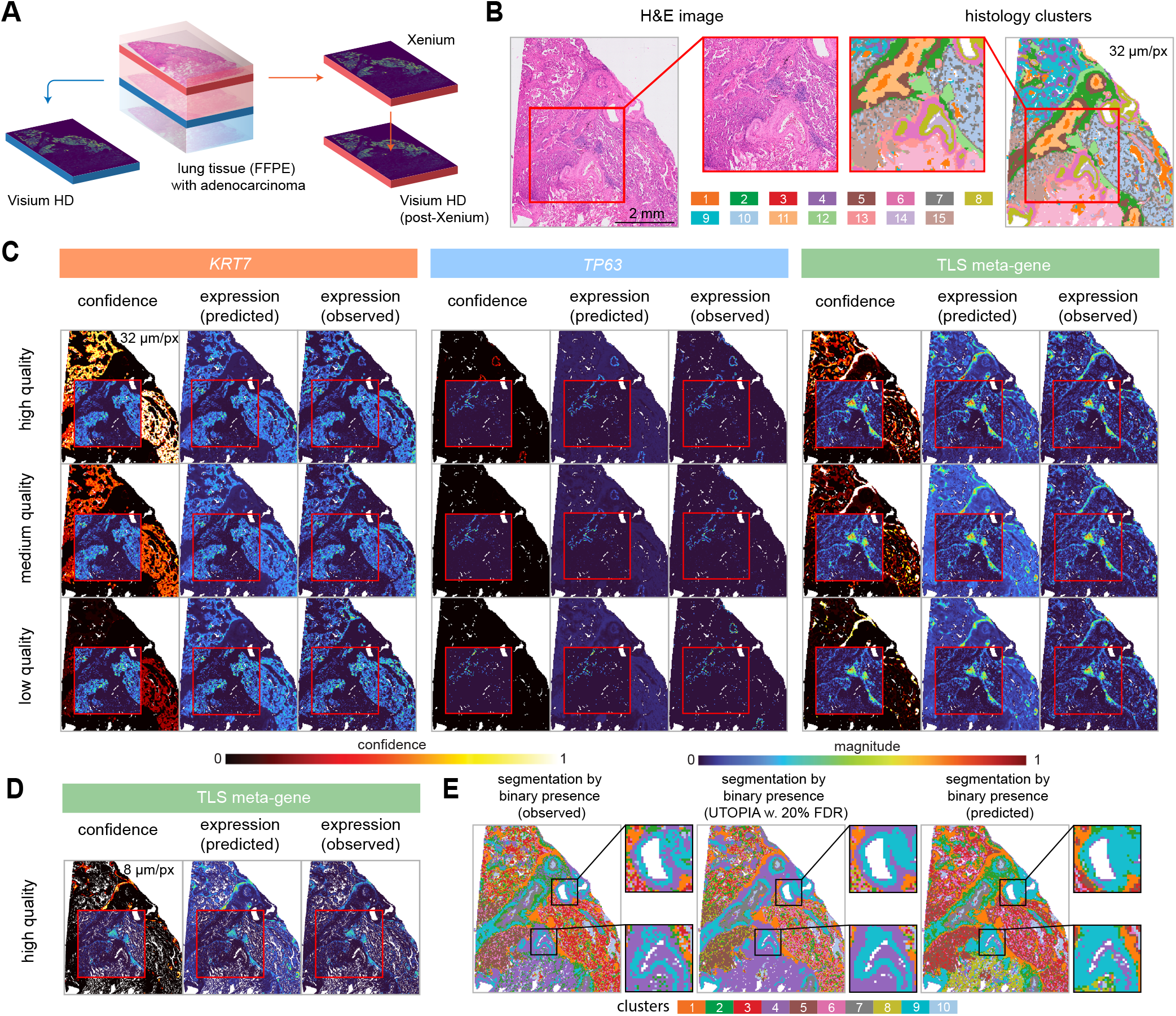
Quality of training data affects confidence scores in UTOPIA. **A**, The setup of measurements by Xenium and Visium HD on two slides in the same tissue block. **B**, The H&E image and histology clusters (32 µm/pixel) of the area with all three measurements covered. One ROI of size 3.2 mm by 3.2 mm was selected. **C**, First column: confidence scores on the existence of *KRT7, TP63*, and TLS meta-gene at 32 µm/pixel. Second and third columns: predicted and ground-truth magnitudes at 32 µm/pixel. Ground-truth magnitudes are displayed inside the ROI. **D**, Performance of model trained on high-quality data at 8 µm/pixel for TLS meta-gene. **E**, Segmentation results by binary gene expression, where the prediction model was trained on high-quality data at 32 µm/pixel. For UTOPIA, we took confidence scores higher than 0.52 (corresponding FDR 20%) as positive expression. For ground-truth and prediction, we took magnitudes greater than 0.05 as positive.

We selected a 3.2 mm × 3.2 mm ROI containing representative histology clusters from the whole slide (**Fig. 6B**). Following the in-sample virtual prediction workflow, we trained three separate prediction models using the three datasets of varying quality restricted to the ROI and then predicted gene expression across the unmeasured regions. UTOPIA was applied to each set of predictions to quantify confidence for the presence of individual genes and meta-genes (see **Methods**). This setup enables a direct comparison of how training data quality affects confidence scores while maintaining a constant prediction target.

The model trained on higher-quality data consistently yielded higher confidence scores than those trained on medium or low-quality data at the 32 μm resolution (**Fig. 6C**). This difference stems from the calibration process: UTOPIA compares predictions against ground-truth measurements within the ROI, and low-quality ground-truth data exhibit sparse and noisy patterns even at coarse resolution. When ground truth appears sparse but predictions exhibit smooth spatial patterns, UTOPIA assigns low confidence, reflecting the mismatch between predicted smoothness and observed sparsity. UTOPIA can thus be conservative when the measured ground-truth is noisy. Therefore, high-quality ground truth with robust molecule counts is essential for generating high-confidence predictions.

Current ST technologies face fundamental limitations: no platform can capture the complete biological truth at single-cell or 8 μm resolution due to technical constraints and the stochastic nature of mRNA expression. However, aggregating measurements to coarser spatial resolutions (e.g., 32 μm per pixel) substantially improves signal quality. Similarly, meta-genes, which are composed of multiple functionally related genes, show markedly improved measurement quality compared with individual genes. For the TLS meta-gene, even the low-quality model achieved high confidence for presence detection at the 32 μm resolution, while the high-quality model attained high confidence even at the finest 8 μm resolution (**Fig. 6D**).

The confidence scores further translate into improved performance on downstream biological analyses. Previous research ^29^ has demonstrated that binarized gene expression (presence/absence) is sufficient for spatial clustering. At a spatial resolution of 32 μm per pixel, binary gene expression derived from UTOPIA’s confidence scores yielded clustering results that closely matched the ground-truth clustering, while exhibiting reduced noise compared to the true binary expression (**Fig. 6E**). In contrast, binary expression obtained from naive magnitude thresholding produced markedly different clustering patterns. These results confirm that UTOPIA’s principled uncertainty quantification enables more reliable biological interpretation than raw prediction values alone.

Collectively, these results establish that data quality is the primary determinant of achievable confidence levels and demonstrate that UTOPIA successfully translates data quality into calibrated, actionable confidence scores that control false discoveries and improve downstream analysis.

## Discussion

Virtual ST has emerged as a promising strategy to extend molecular profiling beyond the current ST technologies, enabling predictions across unmeasured regions within samples and even across unprofiled samples. However, the lack of principled uncertainty quantification has constrained the interpretability and reliability of virtual ST results. Here, we present UTOPIA, a model-agnostic framework that quantifies prediction confidence for virtual ST across two dimensions: spatial resolution and biological granularity. We demonstrated that UTOPIA’s statistically calibrated confidence scores address the four fundamental questions for virtual ST: which molecular targets can be confidently predicted, at what spatial resolution predictions become trustworthy, at what expression magnitude predictions meaningfully distinguish signal from noise, and where within a tissue predictions can be trusted.

UTOPIA conveys a key message that prediction reliability is not a fixed property of a virtual ST model but depends critically on the biological granularity of predicted targets and the spatial resolution of predictions. Across our experiments, UTOPIA revealed a consistent pattern that predictions at coarser resolutions for aggregated biological structures, such as meta genes or broadly defined cell classes, prove more reliable than those at finer resolutions for individual genes or cell types. Moreover, prediction reliability is fundamentally data-dependent; a model performing well on one dataset may fail on another. For instance, our model cannot reliably predict the single gene *CD4* in the gastric cancer sample, yet achieved high confidence for the single gene *KRT7* in the high-quality lung cancer dataset. Benchmarking studies have revealed uneven model performance across existing datasets ^28^, and thus, uncertainty quantification is necessary. UTOPIA addresses this gap and guides practitioners in navigating prediction uncertainty for their specific analysis.

UTOPIA enables users to select the appropriate spatial resolution and prediction target at which predictions are both statistically supported and biologically interpretable. Although virtual ST models often strive to achieve single-cell or 8 μm super-resolution outputs for every gene or cell type, such predictions frequently deviate from ground truth even when spatial patterns appear visually similar. Both UTOPIA’s confidence scores and direct comparisons between predictions and ground truth confirm this limitation. Importantly, many biological applications, such as detecting TLS or glomerular structures, do not require predictions at the finest resolutions or for individual molecular components. Pathway-level meta-genes or aggregated cell types often suffice for biological interpretation, and resolutions of 32 μm per pixel remain visually informative for identifying spatial patterns. Guided by UTOPIA’s confidence scores, researchers can draw biological conclusions at scales where predictions are reliable while avoiding interpretations at scales where confidence is low.

Beyond in-sample predictions, UTOPIA provides false positive control for out-of-subject virtual ST where ground-truth measurements are unavailable. Our analysis of normal gastric tissue samples demonstrates that visually plausible predictions can introduce biologically misleading signals that propagate into downstream analyses, such as spatial domain annotation. For example, predicted *TFF2* expression in gastric glands could be misinterpreted as early metaplastic changes when, in fact, it represents a prediction artifact. UTOPIA distinguishes genuine biological signals from such artifacts, preventing erroneous biological and clinical interpretations. This capability is particularly critical for exploratory analyses and translational research settings where ground-truth molecular validation is infeasible.

Several limitations of the current framework warrant discussion. First, UTOPIA requires representative calibration data, either from ROIs within the same tissue section or from matched samples in out-of-subject settings. Predictions in tissue regions or disease states poorly represented in calibration data may yield unreliable confidence estimates. For in-sample predictions, careful experimental design with representative ROI selection can address this challenge ^20^. For out-of-subject predictions, building large and diverse ST reference databases will be essential. Second, UTOPIA assumes that histologically similar regions share similar molecular profiles. The assumption may be violated when morphology is weakly coupled to molecular state. Third, while UTOPIA supports flexible threshold-based inference, selecting biologically meaningful thresholds remains application-dependent and may require domain expertise.

Despite these limitations, UTOPIA provides a principled foundation for confidence quantification in virtual ST. As virtual ST models, and spatial multiomics approaches^38, 39^, continue to proliferate, frameworks like UTOPIA become essential for revealing which predictions merit trust. By systematically determining predictability across molecular targets, spatial scales, and tissue contexts, UTOPIA establishes a general blueprint for uncertainty-aware inference in spatial omics, transforming virtual predictions from computational outputs into statistically grounded tools for biological discovery.

## Methods

### Overview of conformal inference

Conformal inference is a non-parametric statistical framework for quantifying uncertainty in predictions, which has been previously utilized to quantify gene expression prediction uncertainty^40^. It requires two types of data: calibration data, for which both the ground-truth value 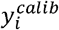 and the predicted value 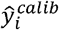 are available, and test data, for which only prediction 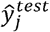 is observed and the goal is to assess whether the corresponding unobserved ground-truth 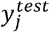 is close to 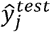.

To illustrate, consider a null hypothesis that the unobserved 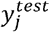 is less than or equal to a constant *c*. Under this hypothesis, the prediction 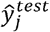 should be consistent with the distribution of 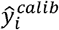 in the calibration data whose ground-truth 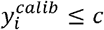. If the prediction 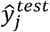 falls outside this calibration distribution, it is deemed non-conformal with respect to the null hypothesis, yielding a small p-value and thus evidence to reject the null. We describe the details in subsequent sections.

### Histology feature extraction and segmentation

We rescaled histology images to a resolution of 0.5 µm per pixel, such that each 16 × 16-pixel tile corresponds to an 8 × 8 µm region, approximately matching the size of a single cell. These tiles, referred to as superpixels in prior literature ^18-20^, serve as the smallest unit for prediction and inference.

To extract feature representations for each superpixel, we cropped a 224 × 224-pixel patch centered on the tile and passed it through a Vision Transformer (ViT)-based pathology foundation model (PFM). The ViT partitions the 224 × 224-pixel patch into a 14 × 14 grid of non-overlapping 16 × 16-pixel tokens. From the model output, we derived two complementary features: a global feature 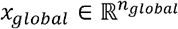, taken from the CLS token, which aggregates information across the entire patch and captures the histological context surrounding the superpixel; and a local feature 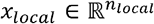, taken from the patch token corresponding to the central 16 × 16-pixel tile, which encodes cell-level morphological detail. Naturally, we have extracted histology features for the H&E image at 8 µm per pixel. For histology features at lower resolutions 8*r* µm per pixel for *r* = 2,3,…, we take the mean of histology features among the *r*^2^ 8-µm bins inside the 8*r*-µm bin. For each resolution, we then applied principal component analysis (PCA) to reduce the dimensionality of the global and local features separately, compressing each to 100 dimensions, and then concatenated them to form a 200-dimensional feature vector. In our implementation, we used UNI as the PFM, which makes *n*_*global*_ and *n*_*local*_ both equal to 1024 prior to PCA reduction.

To identify regions with similar histological characteristics, we performed clustering at each resolution level. We first applied PCA to reduce the 200-dimensional feature vectors to 80 dimensions and then used the K-means algorithm to cluster them into distinct groups.

### Tissue filtering using HistoSweep

We used HistoSweep ^41^ to generate tissue masks for each slide. All subsequent segmentation, training, prediction, and inference by UTOPIA were restricted to pixels within the tissue regions, excluding background areas that appear white on H&E images but contain no tissue.

### Calibration and test data

UTOPIA requires inputs of the calibration data 𝒞 from the calibration area and test data 𝒯 from the test area. The calibration area is the part in which we know the ground truth of gene expression magnitudes or cell type annotations. The test area is the area in which we do not know the molecular profiles, and therefore, we make virtual predictions. UTOPIA infers confidence scores for these predictions. In the case of in-sample broadcasting, the calibration data 𝒞 are the data inside the ROI and the test data 𝒯 are the data everywhere but the ROI.

In particular, we let 𝒞 be a set of (***x***_*i*_,***ŷ***_*i*_,***y***_*i*_) denoting the data for each spot *i* inside the calibration area. Here, the vector ***x***_*i*_ ∈ ℝ^2048^ is the histology feature of spot *i*. The vector 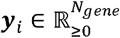 is the ground-truth gene expression magnitudes for *N*_*gene*_ genes in spot *i*. The vector 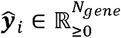 is the predicted gene expression magnitudes for *N*_*gene*_ genes in spot *i* by model *f*. Similarly we write the test data 𝒯 as a set of (***x***_*j*_,***ŷ***_*j*_,***y***_*j*_). The only difference here is that we do not observe the ground truth ***y***_*j*_ for spot *j*.

If the model *f* is predicting cell types instead of gene expression magnitudes, the vector 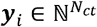 denotes the ground-truth counts of cell types for *N*_*ct*_ cell types in spot *i*. The vector 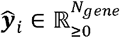 is the softmax scores for *N*_*ct*_ cell types in spot *i*. The same interpretation applies to 𝒯=(***x***_*j*_,***ŷ***_*j*_,***y***_*j*_).

To ease the notation, we stack 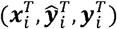 vertically for |𝒞| spots in the calibration data 𝒞 and write 𝒞 =(***X***^*calib*^, ***Ŷ*** ^*calib*^,***Y***^*calib*^).Similarly, we write 𝒯=(***X***^*test*^, ***Ŷ*** ^*test*^,***Y*** ^*test*^). If there is no need of distinguishing calibration and test data, we drop the superscript and use (***x***,***Ŷ***,***Y***). The matrix notation makes it easier to refer the spot information, e.g., ***X***_*i*_,=***x***_*i*_, and the per-gene or per-feature information, e.g., ***Y***_*·,g*_.

### Building predictions in calibration area

In many cases, the calibration data 𝒞 containing the ground truth is the training data for the model *f*. For example, in-sample prediction directly uses the measured ST data in ROI(s) to train a model *f*. Therefore, directly applying the trained model *f* on ***x***^*calib*^ to get,***Ŷ***^*calib*^ risks double-dipping. Instead, we split the calibration data into *N*_*fold*_ folds, where *N*_*fold*_ is data-adaptive and does not require user to specify. Each fold is a rectangle with variable width and height. For the *r*-th fold, we train a surrogate model 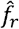 using the same model architecture as the apply the full model *f* by restricting the training data to the calibration data outside the *r*-th fold. We then apply the surrogate model 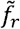. to the *r*-th fold to get the ***Ŷ***^*calib*^ for spots inside the *r*-th fold.

To speed up the process, we need *N*_*fold*_ to be small. On the other hand, to ensure that the surrogate model 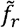 approximate the full model *f*, we cannot have a large fold. Therefore, we develop an adaptive fold-splitting algorithm depending on the histology clustering. Users specify a list of fold sizes, from the largest to the smallest, for UTOPIA to try. By default, fold sizes vary in 1.6 mm × 1.6 mm, 0.8 mm × 0.8 mm, and 0.4 mm × 0.4 mm. UTOPIA tries to split folds using the largest size. If more than 20% spots of any histology cluster inside the ROI(s) fall in one fold, then UTOPIA splits that fold further into smaller sizes until reaching the smallest fold size.

### Conformal inference and FDR control

Although the ground-truth ***Y***^*test*^ is unknown in the test data 𝒯, UTOPIA tells how confident we are such that ***y***_*j,g*_ > *t*_*g*_ for each spot *j* and each gene *g* in the test area, based on available data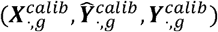 and 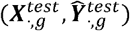. Here the threshold *t*_*g*_ is user-defined.

The null hypothesis for each spot *j* and gene *g* is 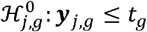 The alternative hypothesis is 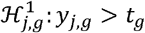. The conformal p-value *p*_*j,g*_ is defined as

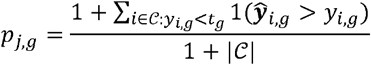

If 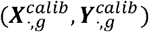 and 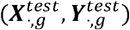 are exchangeable, then applying Benjamini-Hochberg iprocedure on the conformal p-values at level FDR target *q* ∈[0,1]results in a rejection set ℛ such that the expectation of the False Discovery Proportion (FDP) is less than *q*, i.e.,

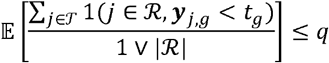

To meet the above regularity condition, we compute the conformal p-values and conduct B-H procedure cluster by cluster. For each histology cluster *k*, define 𝒸_*k*_= {(,***x***_*i*_,***ŷ***_*i*_,***y***_*i*_) ∈ 𝒸: spot, I *i* belongs to cluster *k*} and 𝒯_*k*_ ={(**x**_*j*_,, ***ŷ*** _*j*_,**y**_*j*_) ∈ 𝒯: spot *j* belongs to cluster *k*}. Since similar histology images often yield similar gene expression profiles, we boldly assume that 𝒸_*k*_ and 𝒯_*k*_ are exchangeable. The conformal p-value is thus defined to be

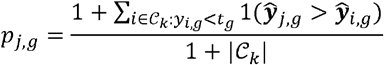

For spot *j* in test data belonging to histology cluster *k*. Afterwards, we apply the B-H procedure for all spots in histology cluster *k* in the test data.

### Null hypothesis in conformal inference

We mostly tested the presence of certain genes or cell types. The threshold for gene is set as 0.05 after normalizing the counts for that gene in the ROIs into the interval [0,1]. resence of a The null hypothesis for each spot *j* and gene *g* is thus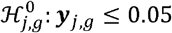. The B-H adjusted conformal p-values indicate the significance of rejecting the null hypothesis and thus how confidence we have for the spot *j* to have expression for gene *g*. The threshold for presence of a cell type is set as 0. Users can choose different thresholds to test in the software.

### Confidence scores

The confidence score for *Y*_*j,g*_ >*t*_*g*_ is defined as the negative log base 10 of the conformal p-value *p* _*j,g*_ after B-H adjustment. In the figures of this manuscript, to improve clarity and contrast,we take the B-H adjusted p-value 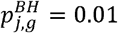as 1 and 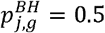as 0. Any value *p*^*BH*^ outside the range is truncated at 0 or 1. Any value *p*^*BH*^ inside the range is normalized in such way

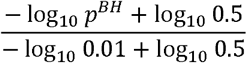

For example, confidence scores at 0.2, 0.5, 0.8 correspond to B-H adjusted p-values at 0.23, 0.07, and 0.02.

### Gene and cell type prediction

Both were predicted using a feed-forward neural network implemented in PyTorch. The model consisted of four fully connected layers with dimensions of 128, 64, 128, and a final output layer corresponding to the number of genes or cell types. Each hidden layer employed ReLU activation followed by dropout equal to 0.1 for regularization. For gene prediction, the final output layer used an exponential linear unit (ELU) activation with parameters α=0.01 and β=0.01 to ensure non-negative predictions suitable for expression values. The model was trained using an asymmetric MSE loss that preferentially penalized underestimation of expression levels. The asymmetric loss penalized two times on underestimation than overestimation. For cell type prediction, we used cross-entropy loss. Both models were trained for 20 epochs with a batch size of 8096 and a learning rate of 0.001. Both models were trained at 8 µm per pixel.

The inputs for both models consist of histology features from 8-µm bins extracted using pathology foundation models, while the supervised signals are the corresponding gene expression values or cell type labels for each bin. For each gene, we normalized the observed counts within regions of interest (ROIs) using min-max normalization to scale values to the interval [0, 1].

### Predictions and ground truth at lower resolution

By default, UTOPIA infers confidence scores on three resolution levels, 32 µm/pixel, 16 µm /pixel, and 8 µm/pixel. The model *f* directly gives a prediction ***ŷ***_*i*_ of resolution at 8 µm/pixel and the ground truth ***y***_*i*_ is also provided at the same resolution. For ***ŷ***_*i*_ and ***y***_*i*_ in lower resolutions, we sum up the predictions and the ground truth of 8 µm bins inside each 16 µm or 32 µm bin. For gene expression, we normalize ***Y***_·,*g*_ to [0,1I] for each gene g and then normalize ***Ŷ*** _·,*g*_ using the same scaling factor. For cell type prediction, we divide the sum of softmax scores by the number of 8-µm bins so that ***Ŷ*** _*·,ct*_ still represents probability; the ground-truth ***Y*** _*·,ct*_is the number of 8-µm bins annotated with the corresponding cell type.

### Meta-gene and aggregated cell type

The ground-truth expression of a meta-gene is defined as the sum of raw counts across all constituent genes, followed by normalization to the interval [0, 1] as described above. For the predicted expression, since individual gene predictions are on the normalized [0, 1] scale, we first rescaled each constituent gene’s prediction back to the original count scale using the inverse of the normalization factor, summed the rescaled predictions, and then renormalized the result to [0, 1] using the same scaling factor as the ground truth.

The ground-truth label of an aggregated cell type is defined as the union of all constituent cell types. Similarly, the predicted probability of an aggregated cell type is computed as the sum of softmax scores across all constituent cell types.

### Joint clustering of H&E images between two normal gastric samples

Since batch effects exist between H&E images from slides N1 and N2, we applied an unpublished in-house feature extraction method to obtain batch-corrected image embeddings. These embeddings were then used for joint clustering via K-means and for model training. The calibration and confidence score inference steps of UTOPIA remain unchanged.

### Cross-fold calibration in the setting of CosMx FOVs

Since CosMx FOVs are small in size but large in number, each FOV naturally serves as a fold in the cross-validation calibration procedure. For each FOV, a surrogate model was trained on data from all remaining FOVs and used to generate predictions on the held-out FOV.

### Detection of glomeruli using connected component analysis

We identified 32 µm pixels with confidence scores exceeding 0.23 (corresponding to 20% FDR) as containing glomerular cell types. Connected components were then extracted using *scipy*.*ndimage*.*label* and filtered by requiring a minimum size of 10 pixels (10,240 µm^2^) and a maximum eccentricity of 0.85 to isolate glomerular structures.

## Data Availability

We analyzed the following datasets in this study: (1) 10x Xenium data from a human patient with gastric cancer, tumor sample (IRB no. 30-2023-1, https://doi.org/10.5281/zenodo.15164980). (2) 10x Xenium data from a human patient with gastric cancer, normal sample (N1) (IRB no. 30-2023-1, https://doi.org/10.5281/zenodo.15164978). (3) 10x Xenium data from a healthy human gastric patient, normal sample (N2) (https://doi.org/10.5281/zenodo.15164947). (4) 10x Xenium 5K data on cervical cancer (https://www.10xgenomics.com/datasets/xenium-prime-ffpe-human-cervical-cancer). (5) 10x Xenium 5K data on ovarian cancer (https://www.10xgenomics.com/datasets/xenium-prime-ffpe-human-ovarian-cancer). (6) CosMx data from human kidney (https://zenodo.org/records/17228449). (7) 10x Xenium and Visium HD data on human lung cancer (https://www.10xgenomics.com/datasets/visium-hd-cytassist-gene-expression-human-lung-cancer-post-xenium-expt).

## Code Availability

UTOPIA was implemented in Python and is publicly available on GitHub at https://github.com/kaitianjin/UTOPIA.

## Acknowledgments

M.L. was partly supported by the following NIH grants R01HG013185, R01LM014592, U19NS135582, R01HL171595, and U01CA294518. L.W. was supported in part by the NIH National Cancer Institute grants U01CA294518, U01CA264583, R01CA266280, U24CA274274, and Break Through Cancer. L.W. is a member of the James P. Allison Institute and the Institute for Data Science in Oncology at The University of Texas MD Anderson Cancer Center and receives research funding from both institutes.

## Author Contributions

This study was conceived of and led by M.L. K.J. designed the model and algorithm with input from M.L., N.Z., Z.R., and Z.C.. K.J. implemented the software and led data analyses. X.Y. and M.Y. provided input on histology image feature extraction. A.S. provided HistoSweep for histology image QC and mask generation. B.D. and K.S. provided CosMx data on kidney. Y.L. and L.W. provided cell type annotations for Xenium data on cervical and ovarian cancer. J.H.P. and T.H.H. provided Xenium data on gastric tissue samples. J.H.P. annotated the gastric cancer sample. K.J. and M.L. wrote the paper with feedback from other co-authors.

## Competing Interests

M.L. receives research funding from Biogen Inc. unrelated to the current manuscript. M.L. is a co-founder of OmicPath AI LLC. T.H.H. is a co-founder of Kure.ai therapeutics and has received consulting fees from IQVIA; these affiliations and financial compensations are unrelated to the current manuscript. L.W. serves as a member of the Scientific Advisory Board for SELLAS Life Sciences and receives compensation outside the scope of this submitted work. The other authors declare no competing financial interests.

## Extended Data Figure Legends

**Extended Data Fig. 1 | Additional information and analysis of the human gastric cancer sample. A**. Left, H&E images of the selected ROIs. Center, histology clusters of the ROIs at 32 µm per pixel (k = 15) with cluster legends in panel B. Right, annotated cell types of the ROIs at 8 µm per pixel, with cell type legends in Fig. 2. **B**. Histology clusters of the whole slide and the two selected ROIs at 16 µm per pixel (top) and 8 µm per pixel (bottom), both with k = 15. **C**. Ground-truth expression of *CD4*, the TLS meta-gene, and TLS cell types at 32 µm per pixel. Colors for *CD4* and the TLS meta-gene represent expression magnitude; colors for TLS cell types indicate cell type presence. **D**. Predicted expression, UTOPIA-inferred confidence scores, and ground truth for *CD4*, the TLS meta-gene, and TLS cell types at 16 µm per pixel. Ground truth was displayed within the ROIs. For TLS cell types, softmax scores were shown as predictions.

**Extended Data Fig. 2 | Additional analysis of the human gastric cancer sample. A**. Predicted expression, UTOPIA-inferred confidence scores, and ground truth for *CD4*, the TLS meta-gene, and TLS cell types at 8 µm per pixel. **B**. Multi-resolution analysis of zoom-in region 3 across 8, 16, and 32 µm per pixel. For *CD4* and the TLS meta-gene, columns show predicted magnitude, ground-truth magnitude, confidence for presence detection, and binarized ground-truth presence. For TLS cell types, columns show softmax probability scores, confidence for presence, and ground-truth cell type annotation. **C**. Predicted expression, UTOPIA-inferred confidence scores for pathway gene sets (angiogenesis, EMT metastatic potential, and complement system), and ground truth at 32 µm per pixel. Ground truth was displayed within the ROIs. FDR control curves were shown at the bottom.

**Extended Data Fig. 3 | Additional analysis of the human cervical and ovarian cancer samples. A**. H&E images, histology clusters, and annotated cell types for selected ROIs. Each sample was independently segmented into 15 histology clusters at 32 µm (cervical) or 40 µm (ovarian) per pixel. Cell types were annotated at 8 µm per pixel. **B**. Predicted and ground-truth distributions of CAF subtypes on the cervical cancer sample at 32 µm per pixel. Predictions were shown as continuous softmax scores; ground truth was shown as binary presence of cell types. Ground truth within the ROI was also displayed. **C**. Same as B, but for the ovarian cancer sample at 40 µm per pixel.

**Extended Data Fig. 4 | H&E image and histology clusters of the human kidney sample. A**. H&E image with 50 selected FOVs indicated by red rectangles. FOV indices were shown in blue. **B**. Histology clusters (k = 10) at 32 µm per pixel. **C**. Magnified H&E images of the 50 FOVs (each 0.5 × 0.5 mm). **D**. Magnified histology clusters at the 50 FOVs at 32 µm per pixel. White pixels indicate regions without tissue.

**Extended Data Fig. 5 | Histology clusters of the human kidney sample. A**. Histology clusters (k = 10) at 16 µm per pixel. **B**. Histology clusters (k = 10) at 8 µm per pixel. **C**. Magnified histology clusters at the 50 FOVs at 16 µm per pixel. White pixels indicate regions without tissue. **D**. Magnified histology clusters at the 50 FOVs at 8 µm per pixel. White pixels indicate regions without tissue.

**Extended Data Fig. 6 | Predicted and ground-truth cell types on 50 FOVs in the human kidney sample. A**. Predicted cell types at 8 µm per pixel across 50 FOVs from the model trained on FOVs from both samples. FOV indices were shown in the top-left corner. White pixels indicate regions without tissue. **B**. Ground-truth (annotated) cell types at 8 µm per pixel across 50 FOVs. White pixels indicate regions without tissue; gray pixels indicate tissue regions lacking annotation. **C**. Predicted cell types restricted to annotated pixels at 8 µm per pixel across 50 FOVs. Per-FOV prediction accuracy was shown in red at the bottom.

**Extended Data Fig. 7 | Predicted cell types and confidence scores on the human kidney sample. A**. Confidence scores for cell type presence at 16 µm per pixel. Columns correspond to cell types; rows correspond to models trained on different FOV subsets. **B**. Same as A, at 8 µm per pixel. **C**. Softmax scores at 16 µm per pixel. **D**. Softmax scores at 8 µm per pixel.

**Extended Data Fig. 8 | FDR control curves on the human kidney sample. A**. FDR control curves of UTOPIA confidence scores for glomeruli, endothelial cells, and podocytes on predictions from the three models. For each model and cell type, three FDR control curves were plotted for UTOPIA performance under 32, 16, and 8 µm per pixel.

