## Supplementary material for "Multiscale confidence quantification for virtual spatial transcriptomics with UTOPIA"

**Extended Data Table 1 | Definition of meta-genes and pathway gene sets.**

| Name | Genes | Notes |
|---|---|---|
| TLS | *CD2, CD3D, CD3E, CD4, CD8A, CD19, CD247, CD27, CD28, CD70, CD79A, IL7R, MS4A1, FOXP3, PDCD1* | Tertiary lymphoid structure signature. Includes markers for T cells (CD2, CD3D, CD3E, CD4, CD8A, CD247, CD28), B cells (CD19, CD79A, MS4A1), co-stimulatory and activation molecules (CD27, CD70), regulatory T cells (FOXP3), immune checkpoint (PDCD1), and lymphocyte survival (IL7R). |
| Angiogenesis | *ANGPT2, PECAM1, VWF, EGFL7, CLEC14A, EDN1* | Angiogenesis pathway gene set. Includes angiopoietin signaling (ANGPT2), endothelial cell adhesion and vascular integrity markers (PECAM1, VWF), endothelial-specific regulators of vascular development (EGFL7, CLEC14A), and vasoactive peptide signaling (EDN1). |
| EMT | *SNAI1, VCAN, TNC, MET, PDGFRB* | Epithelial-mesenchymal transition gene set. Includes transcriptional repressor of E-cadherin (SNAI1), extracellular matrix remodeling components (VCAN, TNC), receptor tyrosine kinase promoting cell motility and invasion (MET), and mesenchymal cell surface marker (PDGFRB). |
| Complement | *CFB, C7, CFHR1, CFHR3* | Complement system gene set. Includes alternative pathway component (CFB), terminal complement complex member (C7), and complement factor H-related regulators (CFHR1, CFHR3). |

**Extended Data Fig. 1 | Additional information and analysis of the human gastric cancer sample. A**. Left, H&E images of the selected ROIs. Center, histology clusters of the ROIs at 32 μm per pixel (k = 15) with cluster legends in panel B. Right, annotated cell types of the ROIs at 8 μm per pixel, with cell type legends in Fig. 2. **B**. Histology clusters of the whole slide and the two selected ROIs at 16 μm per pixel (top) and 8 μm per pixel (bottom), both with k = 15. **C**. Ground-truth expression of *CD4*, the TLS meta-gene, and TLS cell types at 32 μm per pixel. Colors for *CD4* and the TLS meta-gene represent expression magnitude; colors for TLS cell types indicate cell type presence. **D**. Predicted expression, UTOPIA-inferred confidence scores, and ground truth for *CD4*, the TLS meta-gene, and TLS cell types at 16 μm per pixel. Ground truth was displayed within the ROIs. For TLS cell types, softmax scores were shown as predictions.

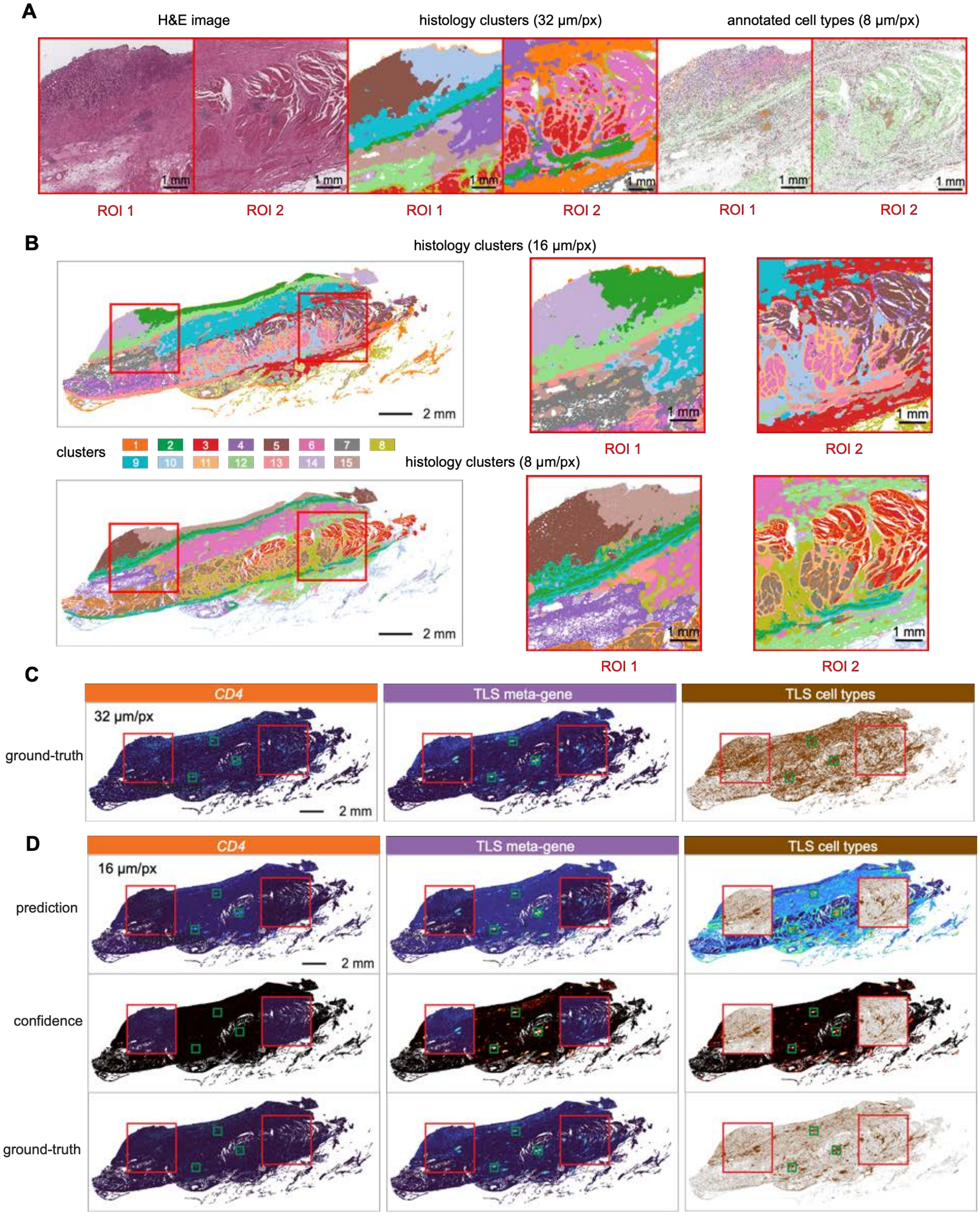

**Extended Data Fig. 2 | Additional analysis of the human gastric cancer sample. A.** Predicted expression, UTOPIA-inferred confidence scores, and ground truth for *CD4*, the TLS meta-gene, and TLS cell types at 8 μm per pixel. **B**. Multi-resolution analysis of zoom-in region 3 across 8, 16, and 32 μm per pixel. For *CD4* and the TLS meta-gene, columns show predicted magnitude, ground-truth magnitude, confidence for presence detection, and binarized ground-truth presence. For TLS cell types, columns show softmax probability scores, confidence for presence, and ground-truth cell type annotation. **C**. Predicted expression, UTOPIA-inferred confidence scores for pathway gene sets (angiogenesis, EMT metastatic potential, and complement system), and ground truth at 32 μm per pixel. Ground truth was displayed within the ROIs. FDR control curves were shown at the bottom.

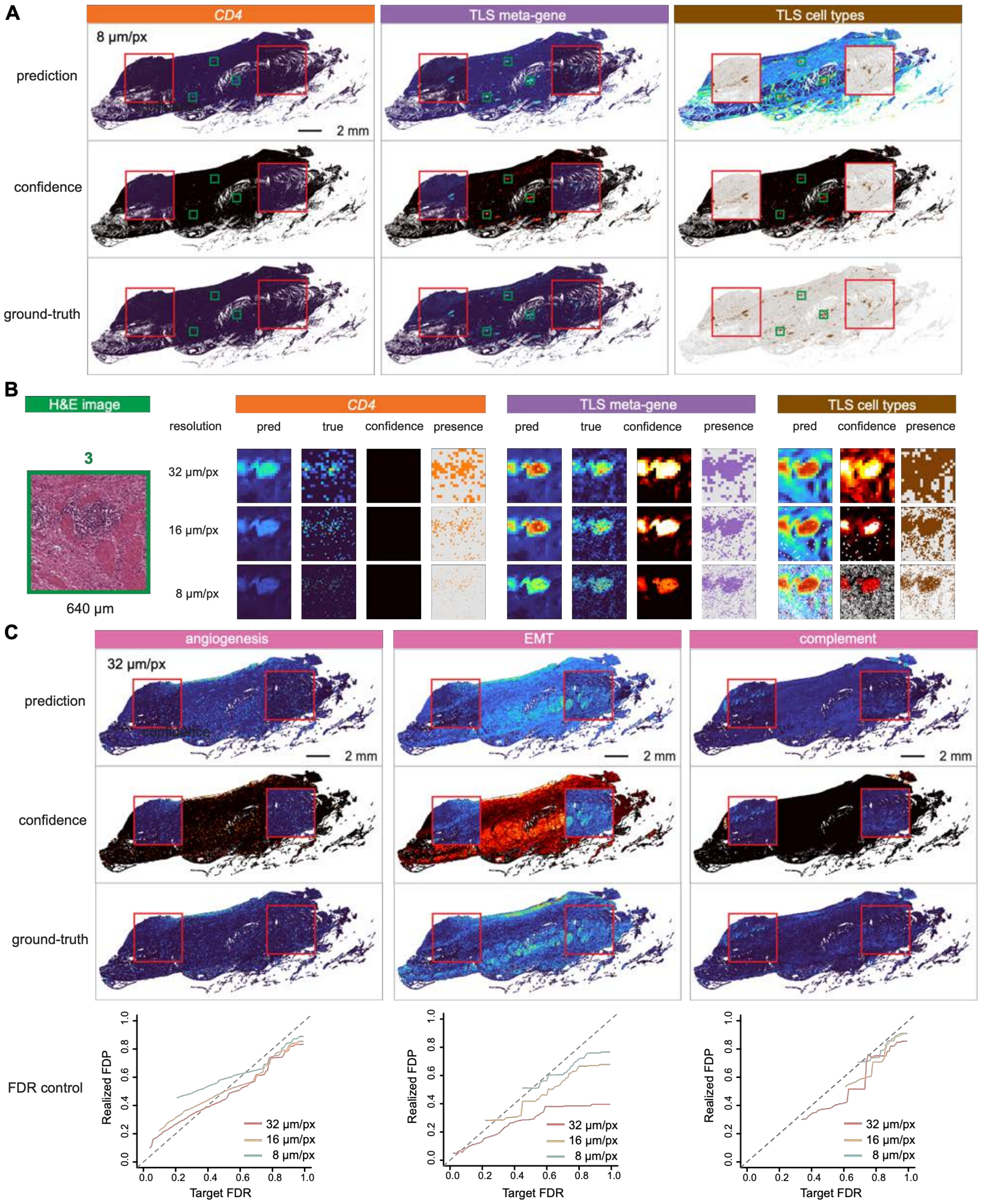

**Extended Data Fig. 3 | Additional analysis of the human cervical and ovarian cancer samples. A**. H&E images, histology clusters, and annotated cell types for selected ROIs. Each sample was independently segmented into 15 histology clusters at 32 μm (cervical) or 40 μm (ovarian) per pixel. Cell types were annotated at 8 μm per pixel. **B**. Predicted and ground-truth distributions of CAF subtypes on the cervical cancer sample at 32 μm per pixel. Predictions were shown as continuous softmax scores; ground truth was shown as binary presence of cell types. Ground truth within the ROI was also displayed. **C**. Same as B, but for the ovarian cancer sample at 40 μm per pixel.

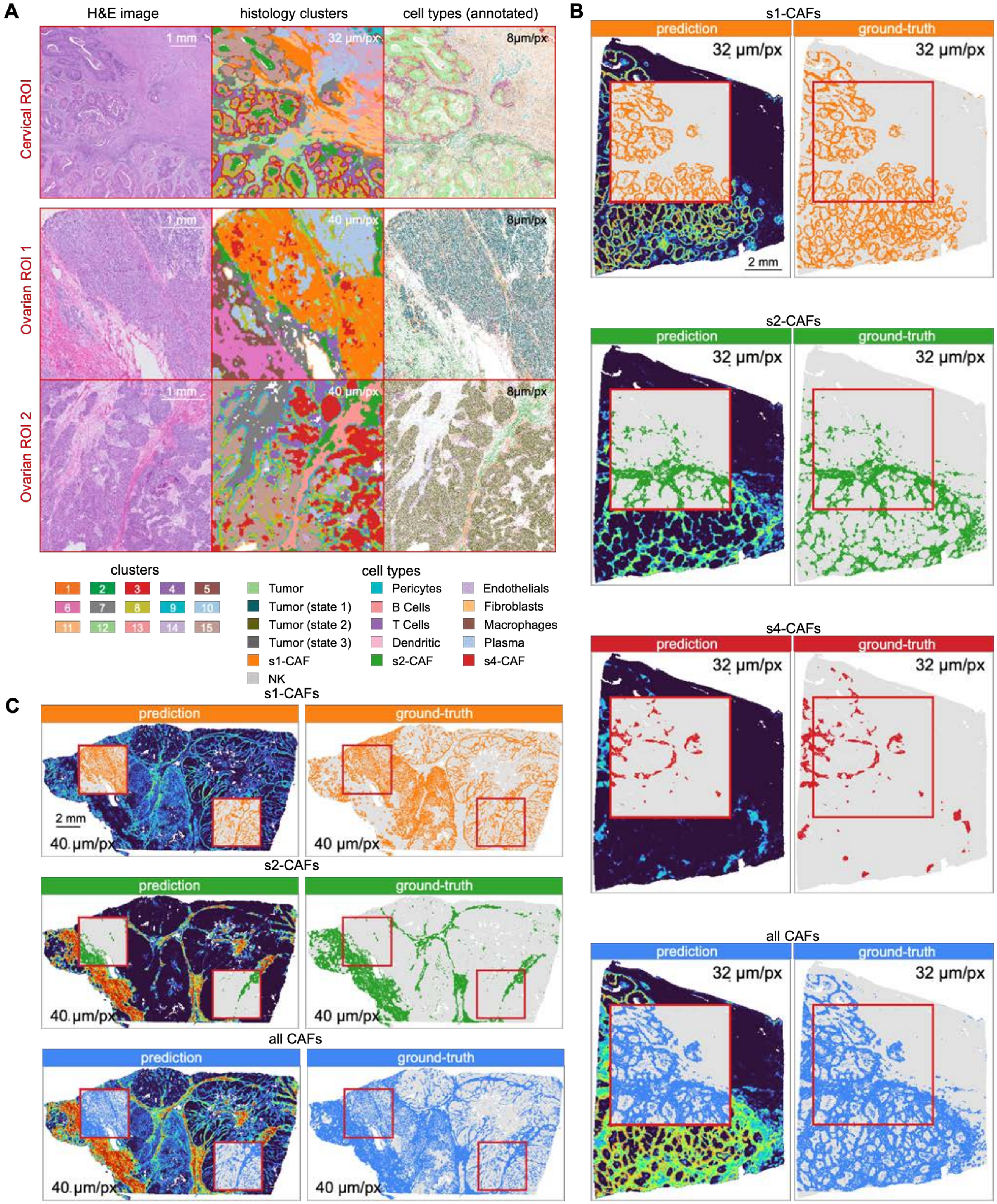

**Extended Data Fig. 4 | H&E image and histology clusters of the human kidney sample. A**. H&E image with 50 selected FOVs indicated by red rectangles. FOV indices were shown in blue. **B**. Histology clusters (k = 10) at 32 µm per pixel. **C**. Magnified H&E images of the 50 FOVs (each 0.5 × 0.5 mm). **D**. Magnified histology clusters at the 50 FOVs at 32 µm per pixel. White pixels indicate regions without tissue.

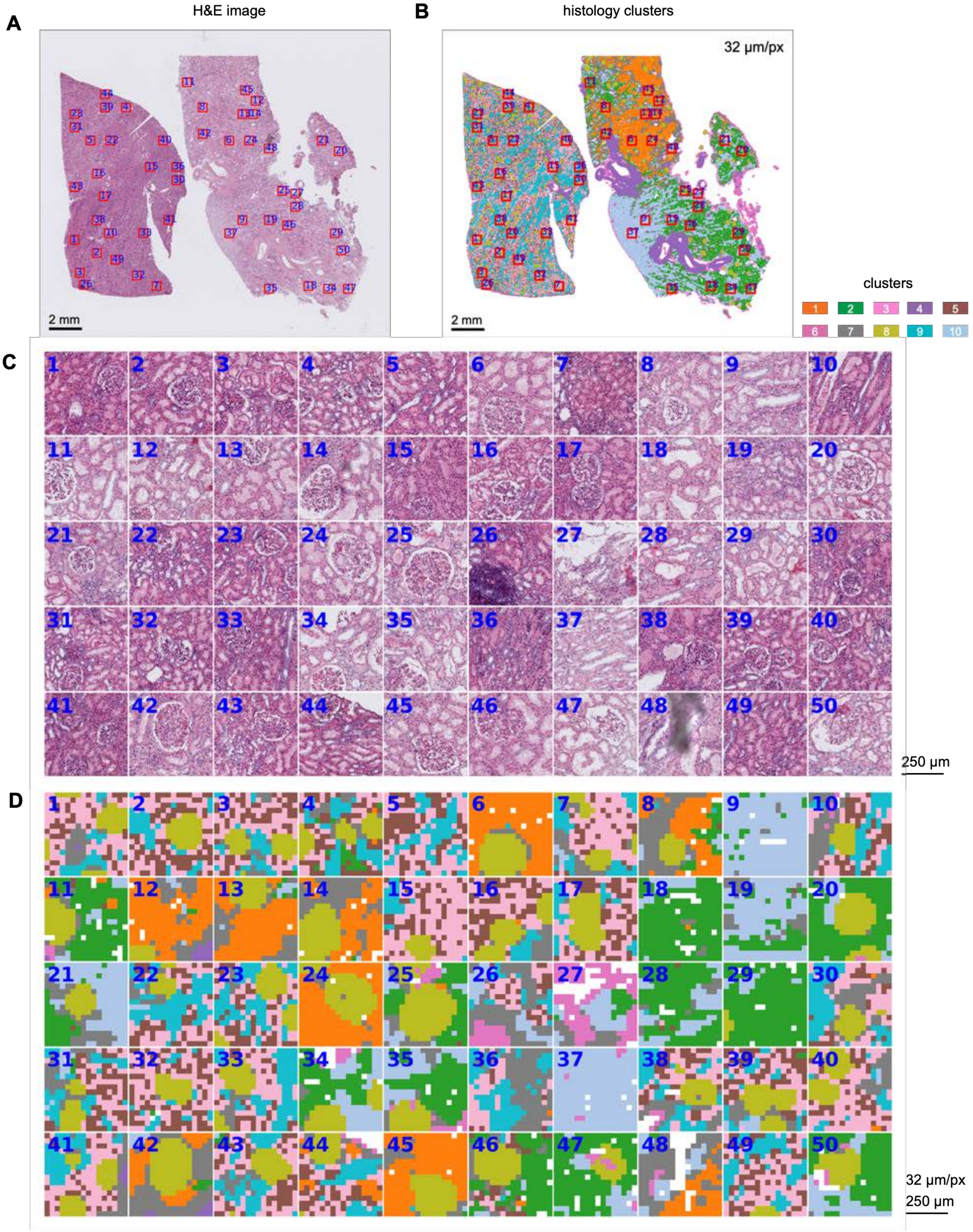

**Extended Data Fig. 5 | Histology clusters of the human kidney sample. A**. Histology clusters (k = 10) at 16 µm per pixel. **B**. Histology clusters (k = 10) at 8 µm per pixel. **C**. Magnified histology clusters at the 50 FOVs at 16 µm per pixel. White pixels indicate regions without tissue. **D**. Magnified histology clusters at the 50 FOVs at 8 µm per pixel. White pixels indicate regions without tissue.

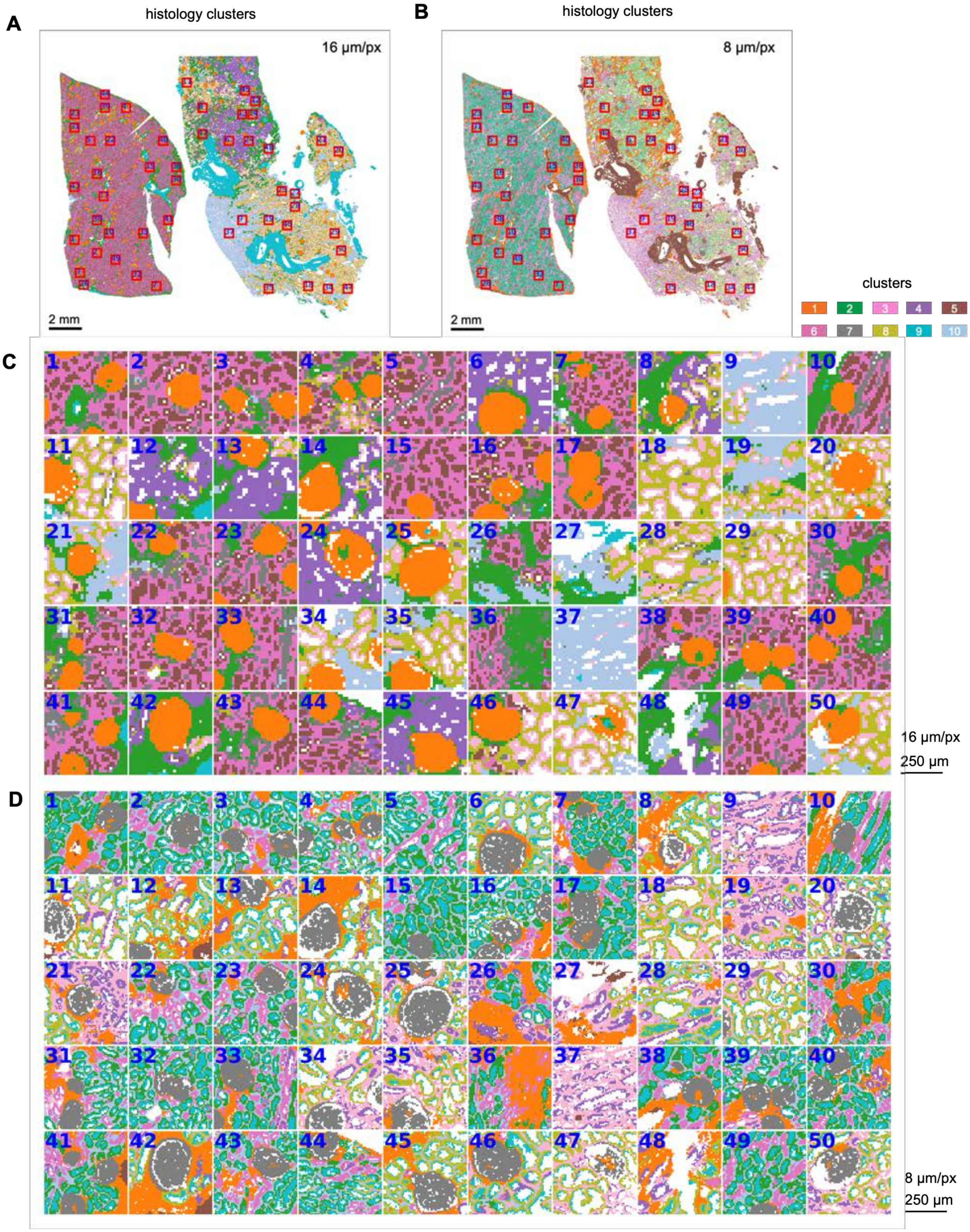

**Extended Data Fig. 6 | Predicted and ground-truth cell types on 50 FOVs in the human kidney sample. A**. Predicted cell types at 8 μm per pixel across 50 FOVs from the model trained on FOVs from both samples. FOV indices were shown in the top-left corner. White pixels indicate regions without tissue. **B**. Ground-truth (annotated) cell types at 8 μm per pixel across 50 FOVs. White pixels indicate regions without tissue; gray pixels indicate tissue regions lacking annotation. **C**. Predicted cell types restricted to annotated pixels at 8 μm per pixel across 50 FOVs. Per-FOV prediction accuracy was shown in red at the bottom.

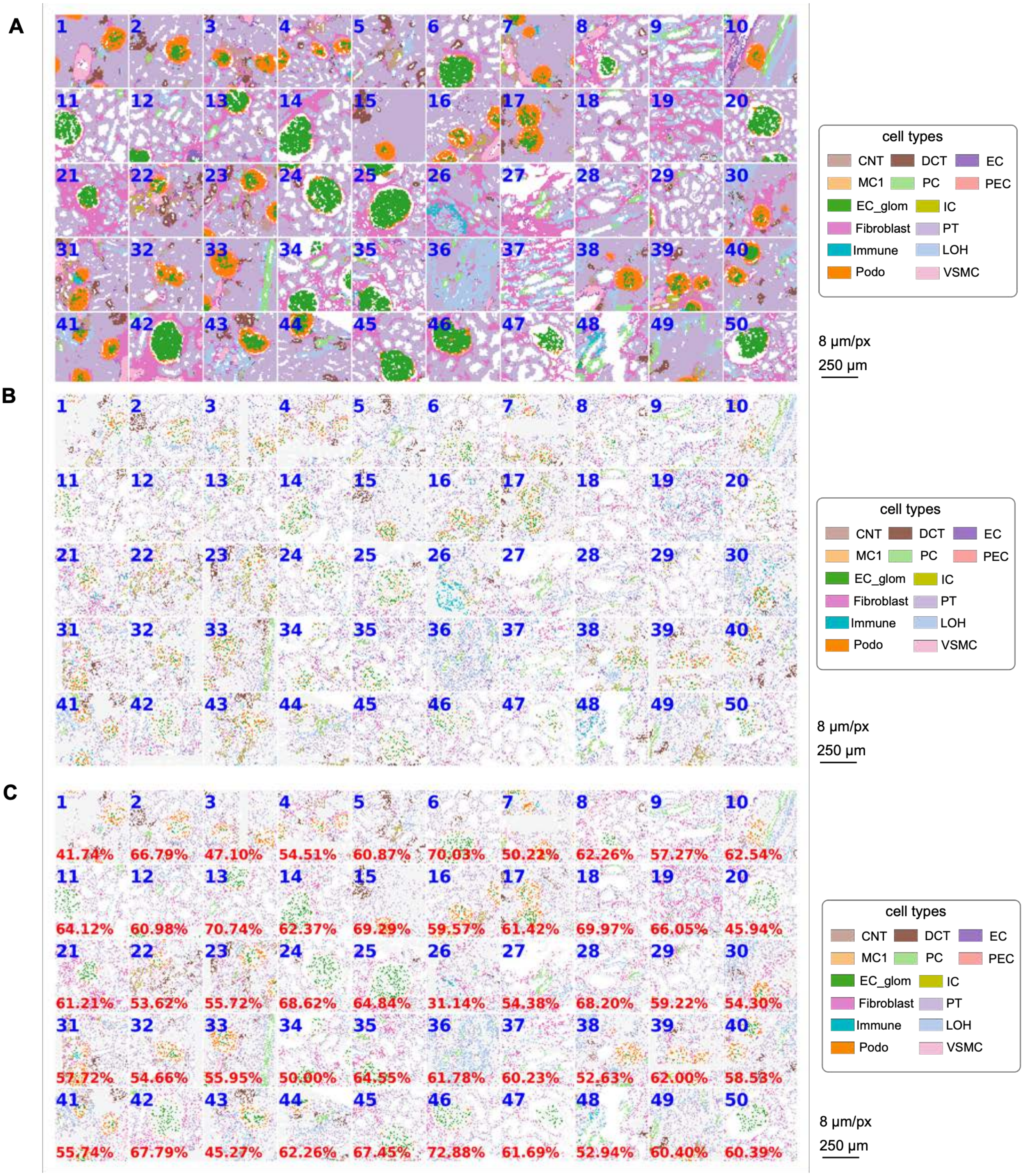

**Extended Data Fig. 7 | Predicted cell types and confidence scores on the human kidney sample. A**. Confidence scores for cell type presence at 16 μm per pixel. Columns correspond to cell types; rows correspond to models trained on different FOV subsets. **B**. Same as A, at 8 μm per pixel. **C**. Softmax scores at 16 μm per pixel. **D**. Softmax scores at 8 μm per pixel.

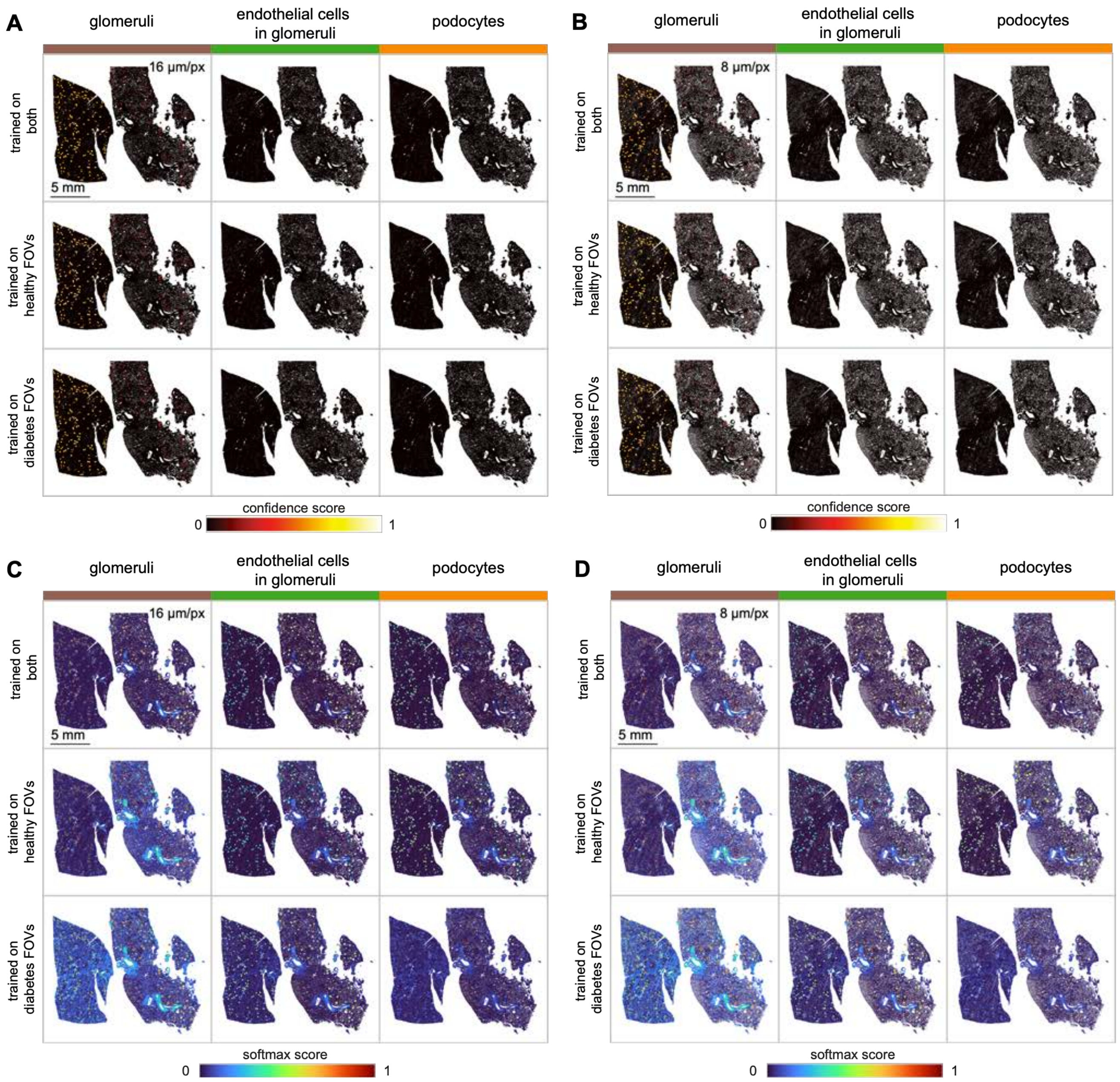

**Extended Data Fig. 8 | FDR control curves on the human kidney sample. A**. FDR control curves of UTOPIA confidence scores for glomeruli, endothelial cells, and podocytes on predictions from the three models. For each model and cell type, three FDR control curves were plotted for UTOPIA performance under 32, 16, and 8 μm per pixel.

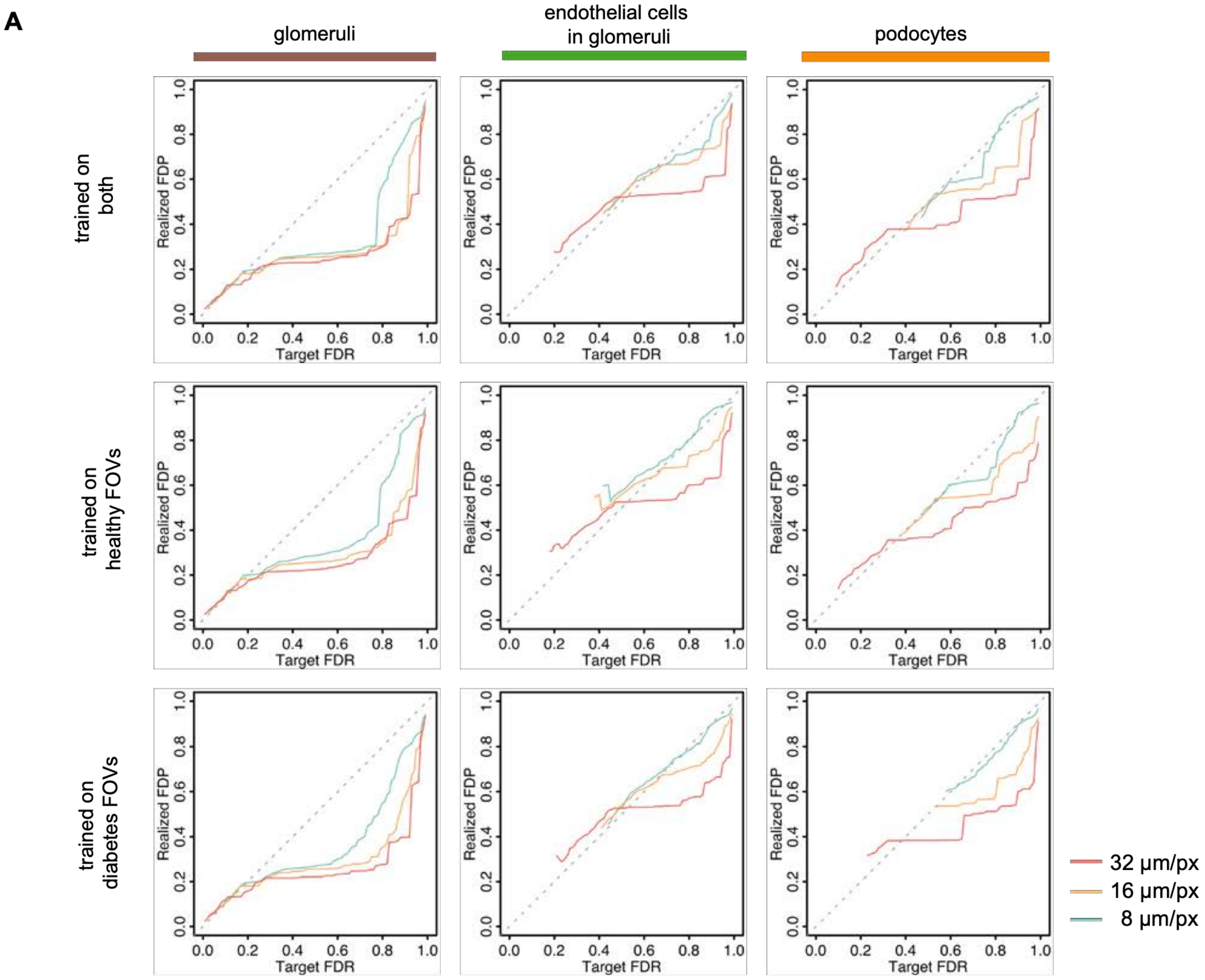
